# Nonlinearities in retinal bipolar cells shape the encoding of artificial and natural stimuli

**DOI:** 10.1101/2020.06.10.144576

**Authors:** Helene Marianne Schreyer, Tim Gollisch

**Affiliations:** Department of Ophthalmology, University Medical Center Göttingen, 37073 Göttingen, Germany; Bernstein Center for Computational Neuroscience Göttingen, 37077 Göttingen, Germany

**Keywords:** Retinal bipolar cells, nonlinearities, contrast representation, spatial integration, LN model, natural stimuli

## Abstract

The retina dissects the visual scene into parallel feature channels, and bipolar cells are speculated to play a key role in this signal separation. Yet, bipolar cells are traditionally viewed as simple, linear neurons. Here, using the salamander retina, we investigated the hypothesis of linear signal processing in bipolar cells by intracellularly recording their voltage signals under artificial and natural visual stimuli. We observed nonlinear representation of contrast and, unexpectedly, also nonlinear spatial integration in a sizable fraction of bipolar cells. Furthermore, linear receptive field models fail to describe responses of nonlinear bipolar cells to spatially structured artificial and natural stimuli. The nonlinear properties occur in the receptive field center and may be cell-type specific, with stronger effects in transient than sustained bipolar cells. Thus, our data suggest that nonlinear signal pooling starts earlier than previously thought, that is, before signal integration in bipolar cells.

## Introduction

Visual processing starts in the eye where the neural network of the retina parses visual signals into dozens of parallel streams of information, represented by different types of retinal ganglion cells (Baden et al., 2016; Gollisch and Meister, 2010; Masland, 2012; Wassle, 2004). A pivotal role in shaping these streams of information is taken up by retinal bipolar cells, which diversify the photoreceptor signals (Euler et al., 2014; Wassle et al., 2009) and provide the basic excitatory input to ganglion cells (Asari and Meister, 2012; Franke et al., 2017). The nature of bipolar cell signals and their transmission to ganglion cells is crucial for how ganglion cells integrate visual information over their receptive fields. For example, the nonlinear spatial integration that is characteristic for Y-type ganglion cells (Enroth-Cugell and Robson, 1966; Krieger et al., 2017; Petrusca et al., 2007) originates in nonlinear inputs to ganglion cells from bipolar cells (Borghuis et al., 2013; Demb et al., 2001; Schwartz et al., 2012). Moreover, nonlinearities in bipolar-to-ganglion cell signaling lie at the root of different computations and feature extractions performed by specific ganglion cells (Baccus et al., 2008; Gollisch and Meister, 2008, 2010; Krishnamoorthy et al., 2017; Munch et al., 2009; Ölveczky et al., 2003; Zhang et al., 2012) and are crucial ingredients in mechanistic models of contrast adaptation (Jarsky et al., 2011; Ozuysal and Baccus, 2012).

Yet, the exact nature of the nonlinear signal transformation along the transmission of visual information through bipolar cells to ganglion cells has remained incompletely understood. Often, it is assumed that nonlinearities arise only at the level of transmitter release from the axon terminals of bipolar cells (Baccus et al., 2008; Borghuis et al., 2013), whereas voltage signals at the soma of bipolar cells are hypothesized to be linearly related to visual contrast (Shapley, 2009). The linear responses of bipolar cells are thought to follow from the linear response characteristics of photoreceptors (Baccus and Meister, 2002; Baylor et al., 1974; Rieke, 2001; Tranchina et al., 1991) and the continual, linear release of neurotransmitter by the photoreceptor ribbon synapse (Heidelberger et al., 2005; Shapley, 2009; Witkovsky et al., 2001).

However, there is little data about how bipolar cells represent complex visual signals in their membrane potential at the level of the soma and whether the presumed linearity actually holds across the diversity of bipolar cell types. Difficulties for studying bipolar cells arise from their relatively inaccessible location in the retina between the layers of photoreceptors and ganglion cells, their small soma size, and their signaling by graded potentials rather than action potentials. This has rendered bipolar cells an inconvenient target for detailed studies of neural coding. So far, the focus has largely been on studying temporal dynamics of bipolar cells through fairly simple light stimuli, such as uniform spots that were flashed or modulated in time (Awatramani and Slaughter, 2000; Baden et al., 2013a; Euler and Masland, 2000; Franke et al., 2017; Werblin and Dowling, 1969). Thus, it remains unclear if and to what degree bipolar cells − at the level of their membrane potential – contribute to nonlinear signal processing in the retina. Moreover, systematic investigations of bipolar cell responses under more complex visual stimuli, such as spatial patterns or natural stimuli, are still lacking and may provide new insight on bipolar cell stimulus encoding.

In this work, we therefore recorded the intracellular membrane potential of bipolar cells in the whole-mount salamander retina under artificial and natural visual stimulation in order to investigate the following questions. (1) Do bipolar cells represent contrast in a nonlinear way? (2) Are there nonlinearities in bipolar cell signal integration? (3) Do nonlinearities affect the encoding of artificial and natural stimuli in bipolar cells? And (4) do nonlinearities occur in specific subsets of bipolar cells?

## Results

To explore potential nonlinear signal processing within retinal bipolar cells, we used sharp microelectrodes to intracellularly record their somatic voltage signals in the whole-mount salamander retina under visual stimulation (Figure 1A). To identify bipolar cells and differentiate them from photoreceptors and amacrine cells, we monitored the recording depth and filled recorded cells with a neuroanatomical tracer to view their morphology after the recording (Figure 1B). In addition, retinas were placed onto a multielectrode array during the recording so that ganglion cell spiking activity could be observed in response to current injection into the recorded cell to verify excitatory effects of putative bipolar cells (Figure 1C, see also Figure S1 and Methods for more details).

**Figure 1.**
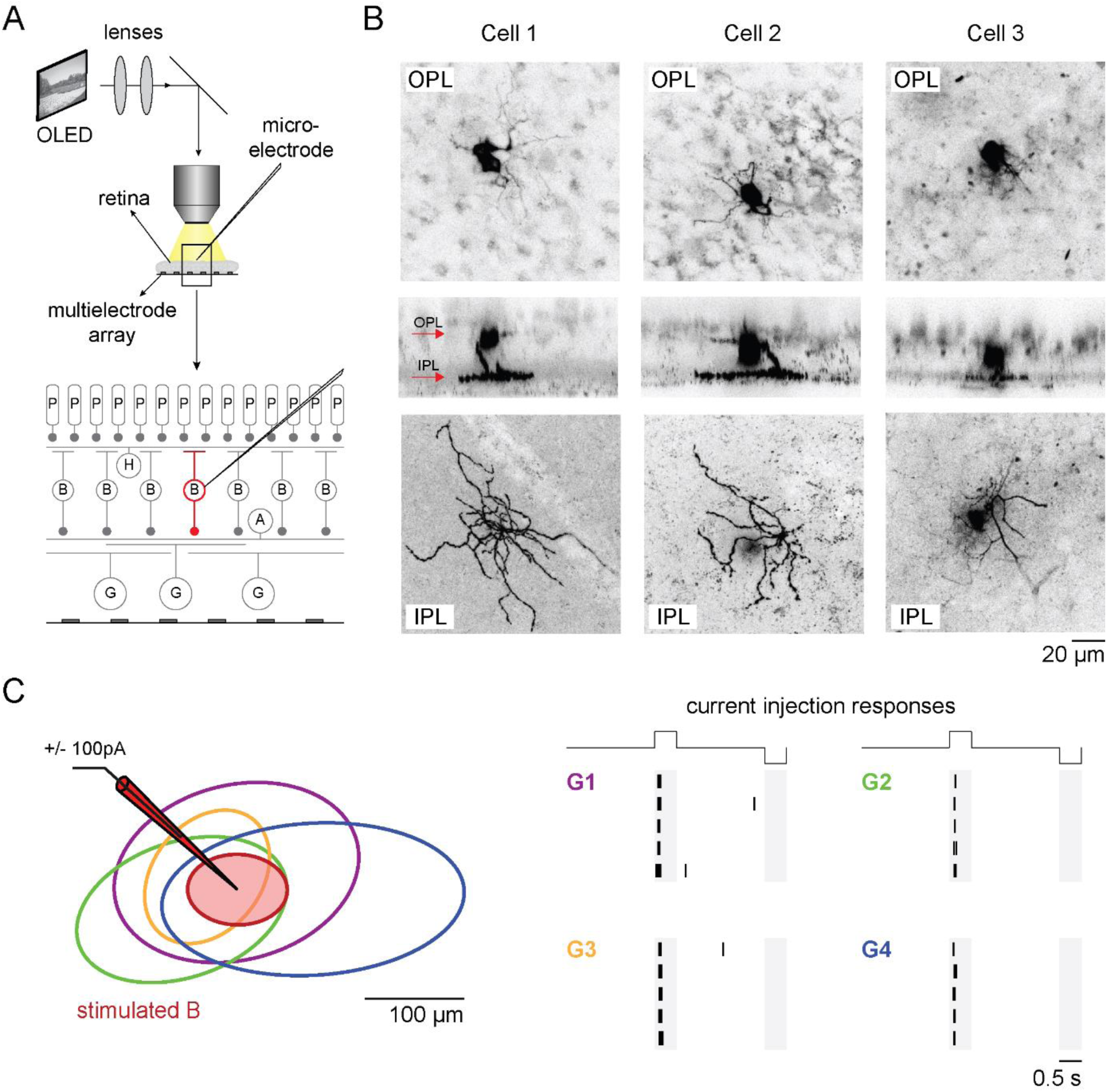
Bipolar cell recordings in the whole-mount retina. A. Schematic view of the experimental setup. The sharp microelectrodes were inserted from the photoreceptor (P) layer to the bipolar cell (B) somata. Ganglion cells (G) faced a perforated multielectrode array. B. Sample morphologies of three recorded bipolar cells. The rows show horizontal sections through the outer plexiform layer (OPL; top) and through the inner plexiform layer (IPL; bottom) as well as vertical sections (center; red arrows mark the OPL and IPL location). The images are maximum projections over manually selected regions. C. Ganglion cell responses to current injection into a bipolar cell. Left: Receptive field outlines of a recorded bipolar cell (red) and four simultaneously recorded ganglion cells. Right: Spike responses of the same four ganglion cells (colors matching to receptive field outlines) to current injection of positive (+100 pA, first gray-shaded region) and negative current (−100 pA, second gray-shaded region) into the bipolar cell (only 5 trials are shown).

### Nonlinear contrast representation in bipolar cells

To test whether bipolar cells represent contrast in a linear or nonlinear fashion, we used spots to stimulate their receptive field centers with steps of increased (“white”, positive contrast) or decreased (“black”, negative contrast) light intensity on a gray background (Figure 2A, left). In order to evaluate membrane potential changes under preferred and non-preferred spot contrast, baseline membrane potentials, as measured during the background illumination prior to spot presentation, were subtracted. Figure 2A shows trial-averaged response traces for three sample cells. Cell 1 responded with similar amounts of depolarization and hyperpolarization to the preferred and non-preferred spot contrast, respectively, consistent with a linear response. Cell 2 had stronger depolarization than hyperpolarization, suggesting a mild nonlinearity; and Cell 3 only depolarized and did not show any hyperpolarization for the non-preferred spot, indicative of strongly nonlinear, completely rectifying response characteristics. To quantify how nonlinearly a cell represented preferred versus non-preferred contrast, we computed a hyperpolarization index (HPi) as the normalized difference between the peak depolarization and the peak hyperpolarization in the two spot response traces (see Methods). The index takes values close to zero for equal amounts of hyper- and depolarization (corresponding to linear responses), values close to unity when there was hardly any hyperpolarization compared to the depolarization (strong nonlinearity), and negative values for larger hyperpolarization than depolarization. We found that the majority of our recorded cells showed evidence of nonlinear representation of preferred versus non-preferred contrast (Figure 2C).

**Figure 2.**
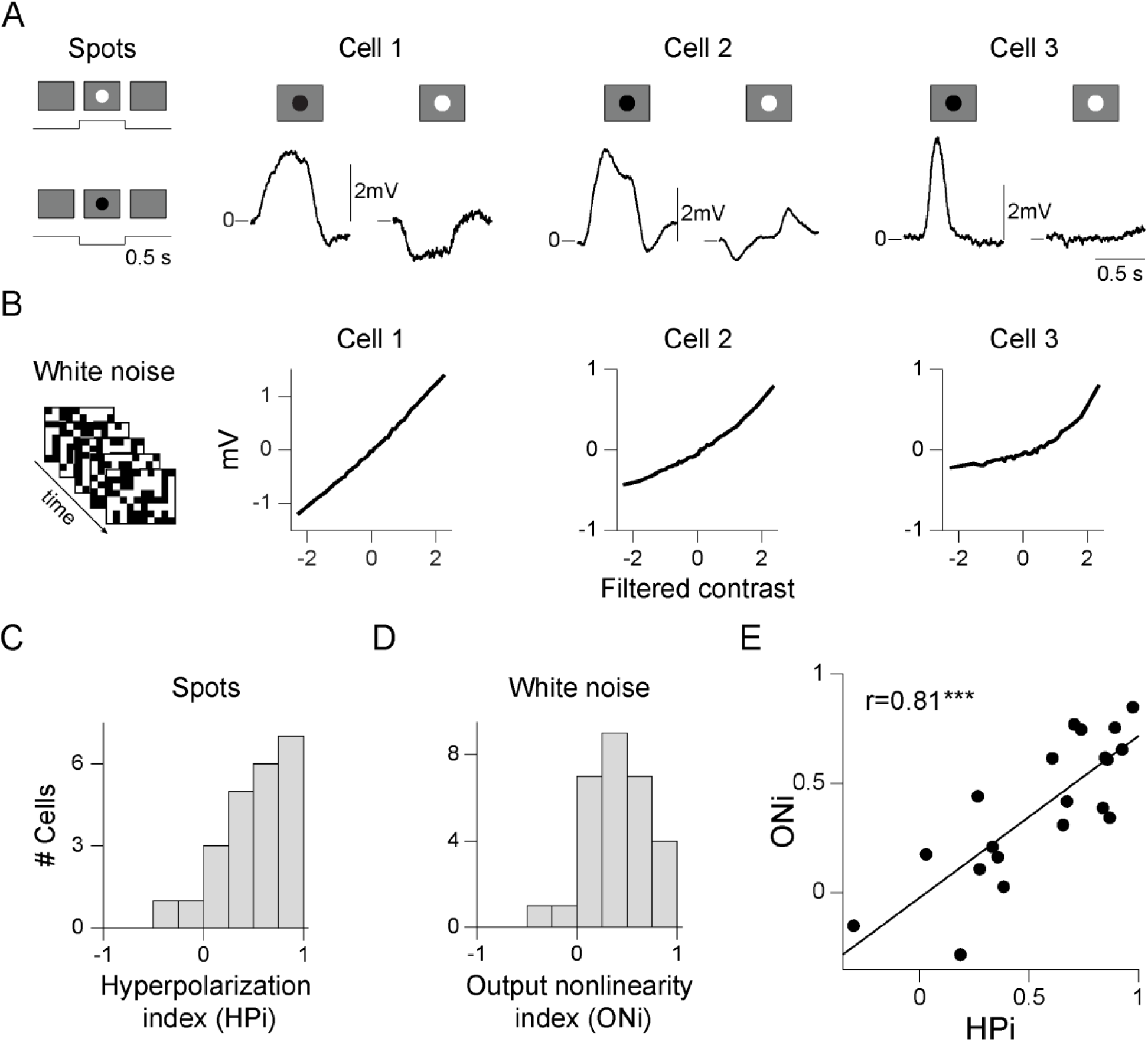
Nonlinear contrast representation in bipolar cells. A. Responses to contrast steps. Left: Schematic display of the spot stimulus with 500-ms contrast steps. Right: Trial-averaged response traces of three sample cells from spot onset to 500 ms after spot offset. Here and in subsequent plots, the zero line marks the baseline membrane potential, measured under background illumination. B. Responses to spatiotemporal white noise. Left: Schematic display of the stimulus, displaying individual frames of binary white noise. Right: Output functions (40 averaged bins) of the same three sample cells as in A. C Distribution of measured hyperpolarization index values (HPi, n=23). D. Distribution of measured output nonlinearity index values (ONi, n=29). E. Comparison of HPi and ONi. Each data point corresponds to a different bipolar cell (n=20). The two indices were significantly correlated (regression line shown as black line, *** p<0.001).

We next asked whether the representation of contrast in bipolar cells changes when switching from contrast steps to continuous, dynamic stimulation. To test this, we applied visual binary white noise on a checkerboard layout and analyzed each recorded cell with a linear-nonlinear (LN) model. The LN model is composed of a linear stimulus filter over space and time, which can be obtained from the spike-triggered average (Chichilnisky, 2001), and a nonlinear transformation, the cell’s output function, which relates the filtered contrast to the membrane potential (see Methods). The shape of this output function directly indicates whether the filtered contrast signal is represented linearly or nonlinearly by the bipolar cell’s membrane potential.

Figure 2B shows the output functions for the same three sample cells, ranging again from linear (Cell 1) over slightly nonlinear (Cell 2) to strongly nonlinear (Cell 3) shapes. We quantified the degree of nonlinearity in the output function by computing an output nonlinearity index (ONi) that compared the average gain (obtained as the slope of a fitted straight line) in the range of preferred (positive filter output) and non-preferred (negative filter output) contrast sequences (see Methods). The index is close to zero for linear output functions, larger than zero for cells that show stronger gain in the depolarization as compared to the hyperpolarization, and can be smaller than zero for cells that show saturation at large filter output. The distribution of obtained ONi values (Figure 2D) again indicated diverse degrees of nonlinearity, ranging from linear to strongly nonlinear.

When comparing the degree of nonlinearity under contrast steps and dynamic white noise stimulation, we observed a pronounced positive correlation (Figure 2E, r=0.81, p=2×10^−5^, n=20). Thus, whether a cell represented contrast linearly or nonlinearly did not depend on the applied stimulus, but appears to be a cell-specific property.

### Nonlinear spatial integration in bipolar cells

Bipolar cells integrate visual information over space by pooling signals across their dendritic branches from several photoreceptors. The mechanisms that generate the nonlinear contrast representation observed above could occur either before this spatial pooling or afterwards. The former would imply that visual signals inside a bipolar cell’s receptive field are not simply summed but nonlinearly combined. To test for this possibility, we applied visual stimuli that subdivided the receptive field center into regions of opposing stimulation, e.g. a bright and a dark half (“split spot”), and periodically reversed the contrast polarity every half second (Figure 3A, top). If the nonlinear mechanisms occur only after input integration, the positive and negative activation from the two opposing contrasts should cancel each other, and the bipolar cell should not respond to the polarity reversals.

**Figure 3.**
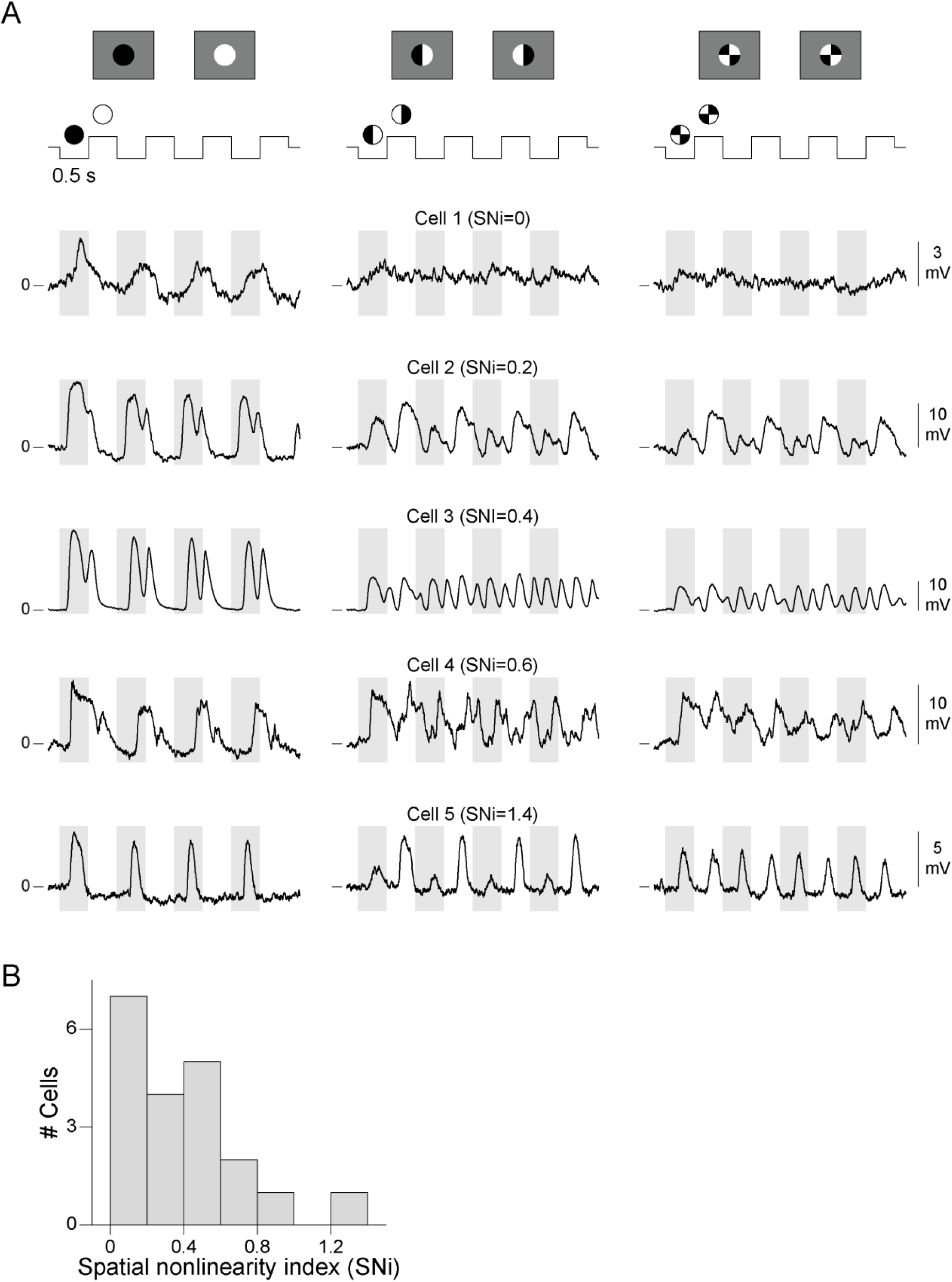
Nonlinear spatial integration in bipolar cells. A. Original single-trial voltage traces of five bipolar cells in response to a contrast-reversing uniform spot (left column), patterned spot divided into two halves (“split spot”, middle column), and patterned spot divided into four quarters (right column). The traces are shown from 200 ms before stimulus onset to 200 ms after stimulus offset. For each cell, the displayed spatial nonlinearity index (SNi) corresponds to the maximum over all spatial patterns. For some cells (e.g. Cell 3), we observed a frequency quadrupling (4 peaks per stimulus period, see also Figure S2B). B. Distribution of the spatial nonlinearity index (n=20).

Figure 3A shows the traces of five bipolar cells to the split spot (middle column) as well as to a uniform contrast-reversing spot for comparison (left column). All five cells clearly responded to the uniform spot (baseline condition). Yet, under reversals of the split spot, Cell 1 showed hardly any deviation from baseline membrane potential, indicative of linear spatial integration, whereas Cells 2-5 responded with depolarization to both reversals. This frequency doubling in the responses of these four sample cells is a telltale sign of nonlinear spatial integration (Demb et al., 1999; Enroth-Cugell and Robson, 1966; Hochstein and Shapley, 1976a, b; Petrusca et al., 2007). To quantify the degree of nonlinear stimulus integration, we computed a spatial nonlinearity index (SNi) by computing a Fourier decomposition of the response traces and comparing the power of the first and of higher harmonics (Hochstein and Shapley, 1976b; Turner and Rieke, 2016, see also Methods and Figure S2). The spatial nonlinearity index (SNi) was close to zero if the cell did not respond to the split spot reversals, whereas values larger than zero indicated frequency doubling and thus nonlinear stimulus integration. As additional “patterned spot” stimuli besides the split spot, we also used a spot divided into four quarters (right panel in Figure 3A), and spots divided into fine checkerboard layouts (see Methods). We approximated the degree of nonlinear stimulus integration per cell by taking the maximum SNi over different spatial patterns (Hochstein and Shapley, 1976b). The observed maximum SNi values (Figure 3B) indicate that bipolar cells covered a range from linear stimulus integration (SNi≈0) to clearly nonlinear integration (up to SNi≈1.4).

### Nonlinear spatial integration curbs the prediction accuracy of the linear-nonlinear model

In order to further analyze how the observed nonlinear properties affect the encoding of visual stimuli, we tested how well bipolar cell responses are captured by the LN model, which had already been used above (Figure 2B) to assess nonlinear contrast representation. The model aims at capturing a cell’s response by first performing a linear integration of the stimulus via a spatiotemporal filter and then passing the result through a nonlinear transformation (capturing, for example, rectification and/or saturation). Thus, the LN model can accommodate nonlinear representations of contrast, but not nonlinear spatial integration. In the retina, the model has often been used to predict responses of ganglion cells to artificial and natural stimuli, and the accuracy of the prediction has served as a way to assess whether the cells indeed integrated spatial or temporal inputs linearly (Chichilnisky, 2001; Freeman et al., 2015; Liu et al., 2017; Pillow et al., 2008; Schwartz et al., 2012). For retinal bipolar cells, on the other hand, only few individual examples of LN model investigations exist, which focused mainly on temporal stimulus structure and showed mostly linear output functions and accurate response prediction (Baccus and Meister, 2002; Baccus et al., 2008; Rieke, 2001).

We first looked at LN models under full-field white noise stimulation without spatial structure. For stimuli with white noise statistics, the linear filter and the nonlinearity of the LN model can be easily obtained with reverse-correlation techniques (Chichilnisky, 2001). Figure 4A shows the obtained model components for a spatially linear and for a spatially nonlinear sample cell, as determined by the patterned spots. The stimulus contained periodically inserted short segments of identical white noise sequences (“frozen noise”), which were excluded from the reverse correlation analysis and which served as a held-out test stimulus (see Methods). The cells responded reliably to the different trials of the frozen-noise sequence (standard deviation across trials and averaged over the frozen-noise duration: 0.36 mV ± 0.15 mV, n=11, mean ± SD). Moreover, the predicted responses obtained from the LN model accurately matched the cells’ averaged responses (Figure 4A). To quantify the similarity between prediction and actual response, we computed the explained variance R^2^ as the squared correlation coefficient between model prediction and averaged response. Across the population of recorded cells, this prediction performance ranged from 68-97% (Figure 4C), which indicates that the LN model accurately predicted responses of bipolar cells to stimuli with no spatial structure.

**Figure 4.**
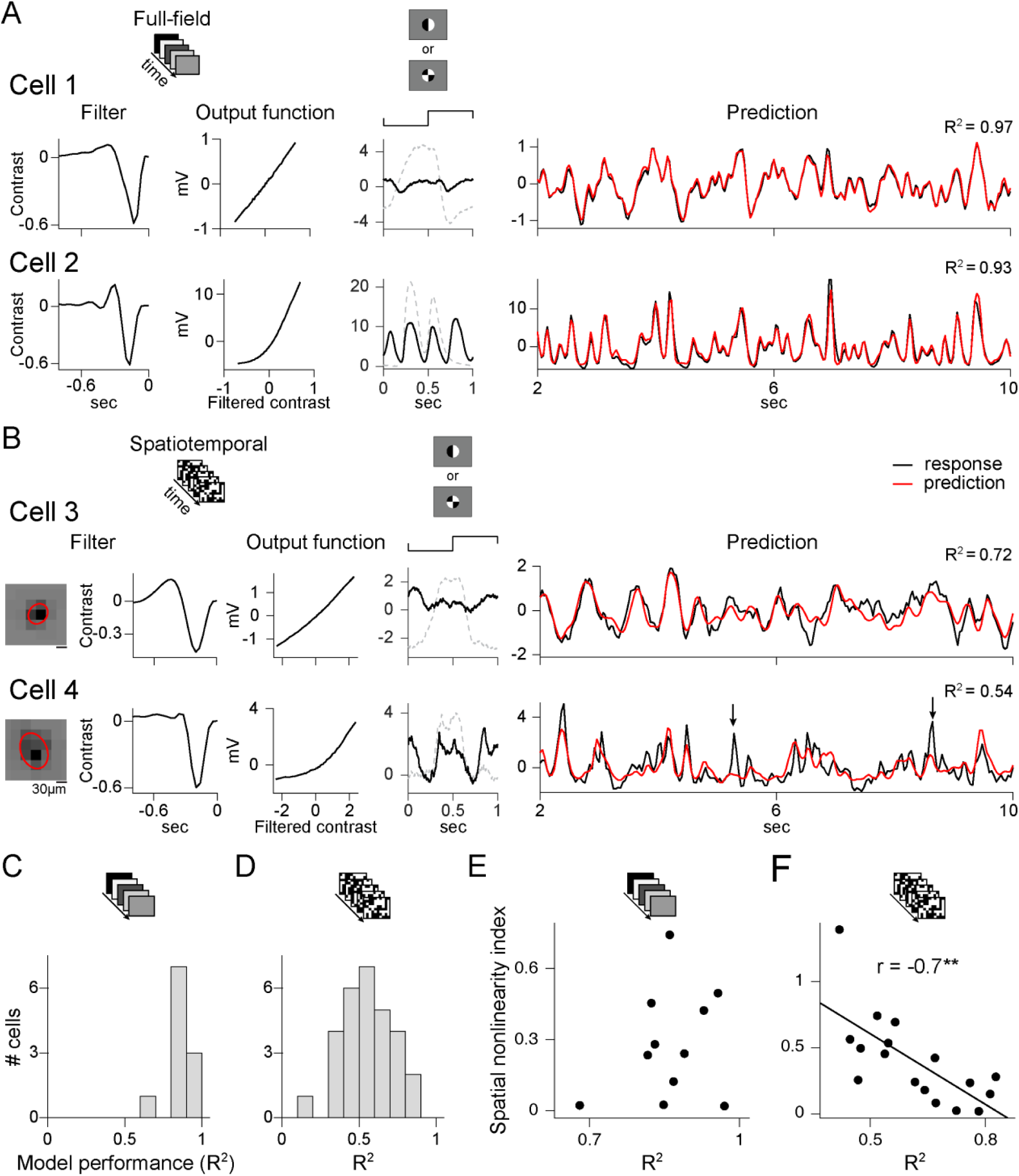
Nonlinear spatial integration curbs the prediction accuracy of the linear-nonlinear (LN) model. A. LN model for full-field temporal flicker for two sample cells. Left: Temporal filters and output functions, with schematic display of the stimulus on top. Middle: Trial-averaged response traces (black line) to the patterned spot for which the frequency doubling was maximal and corresponding trace to the uniform spot (dashed gray line) for comparison. Right: Trial-averaged measured (black) and predicted (red) response traces to a 10-s full-field contrast sequence. B. LN model for spatiotemporal flicker for two sample cells. Left: Spatial and temporal filters as well as the output functions, with schematic display of stimulus on top. Middle: Same as in A for the two sample cells. Right: Measured (black) and predicted (red) response traces to a test stimulus sequence of 10 s. For Cell 4, the predicted and measured response traces match less accurately, and the black arrows mark response peaks that were not predicted by the LN model. C. Distribution of model performance (R^2^) for full-field white noise (n=11). D. Same as C for spatiotemporal white noise (n=29). E. Comparison of the model performance under full-field white noise with the spatial nonlinearity index. Each data point represents one bipolar cell (n=11). There was no significant correlation. F. Same as in E for the spatiotemporal white noise stimulus (linear regression line in black, **p<0.01).

Next, we investigated whether the accuracy of the predictions was affected by spatial structure of the stimulus. To do so, we analyzed bipolar cell responses to spatiotemporal white noise, arranged on a checkerboard with squares of 22.5 or 30 µm. Figure 4B shows the two stages of the LN model for a spatially linear and a spatially nonlinear sample cell. The linear filter is separated into a spatial component (corresponding to the spatial receptive field) and a temporal component. We used the model to predict voltage traces for several held-out segments and assessed model performance by averaging R^2^ over these segments for each cell. The more linear Cell 3 from Figure 4B had fairly accurate predictions with 72% explained variance, whereas the more nonlinear Cell 4 only yielded 54% explained variance. In general, the distribution of the model performance for the spatiotemporal white noise stimulus (Figure 4D) was much broader than that for the spatially uniform white noise stimulus and varied between 13-83% explained variance (n=29). This indicates that the assumed linear integration over space in the model does not correctly describe all bipolar cells.

To test whether indeed the nonlinear spatial integration was causing the low prediction performance of the LN model, we studied the relationship between the spatial nonlinearity index (SNi) and the model performance. For the full-field white noise stimuli, we did not observe a correlation (r=0.24, p=0.47, n=11); spatially linear as well as nonlinear cells showed good model predictions (Figure 4E). For the spatiotemporal white noise stimulus, however, we observed a clear negative correlation (Figure 4F, r=-0.7, p=0.002, n=17); higher spatial nonlinearity indices (i.e., stronger responses to the patterned spots) come with lower performance of the spatiotemporal LN model and vice versa. The results show that the assumed linear integration of the model does not correctly describe all recorded bipolar cells. For example, the LN model would predict activity cancellation and no response when simultaneous opposing contrast occurs inside the receptive field, which may explain why it missed specific response peaks of nonlinear bipolar cells (see arrows in Figure 4B for Cell 4).

### Nonlinear contrast representation and spatial integration under natural movies in bipolar cells

So far, we have considered bipolar cell responses to fairly artificial light stimuli, but natural stimuli have different dynamics and spatial structure. To test whether bipolar cells represent and integrate contrast also nonlinearly under natural stimulus statistics, we recorded responses to natural movies from the “CatCam” database (Betsch et al., 2004), which, although they do not represent the natural habitat of salamanders, contain a variety of natural objects, textures, and motion signals. Cells responded reliably to different trials of a movie (Figure 5A, time-averaged standard deviation 0.34±0.19 mV, n=19 movies, 9 cells).

**Figure 5.**
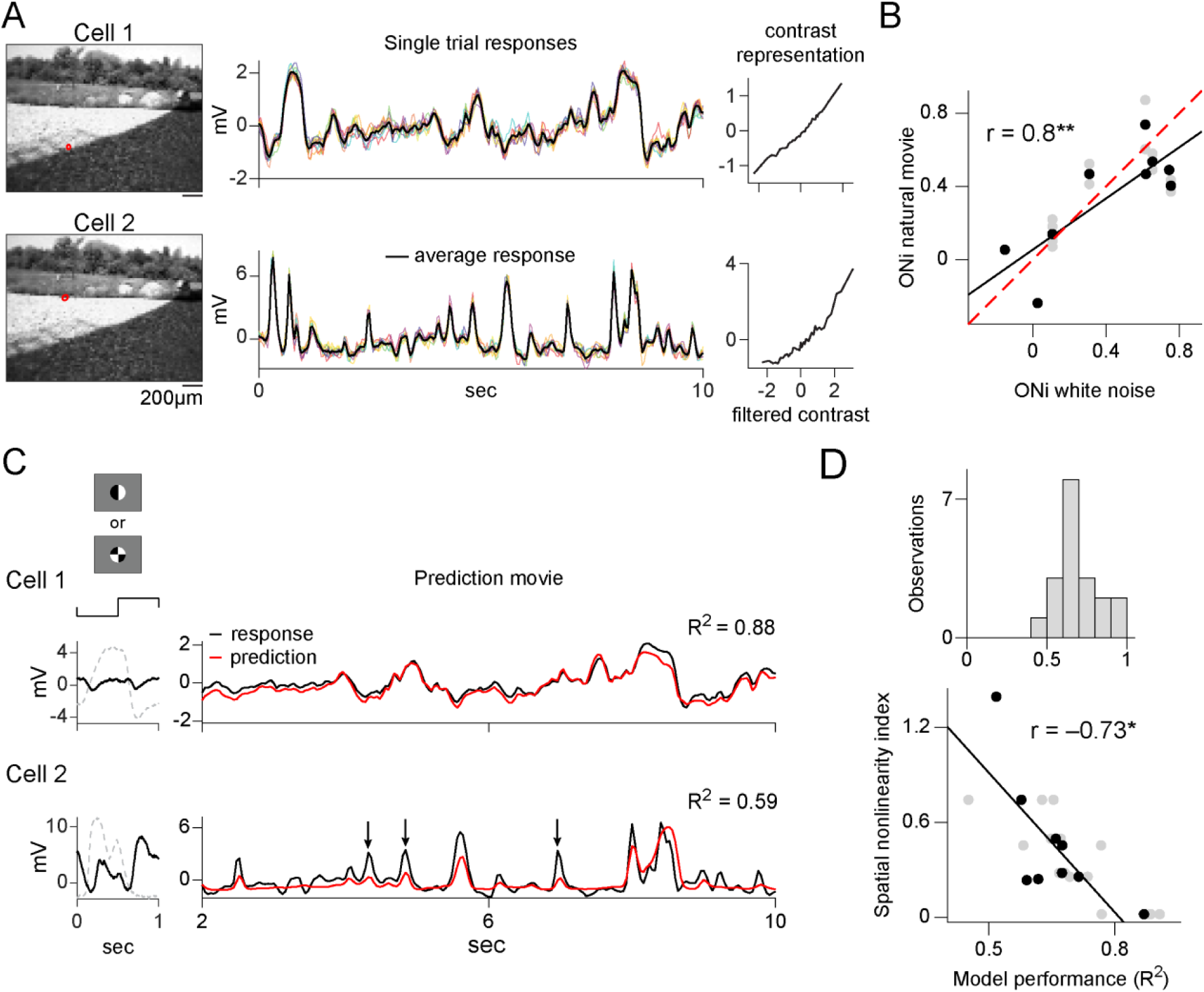
Nonlinear contrast representation and spatial integration under natural movies. A. Responses to natural movies from two sample cells. Left: Single movie frame with receptive field outlines (red circles). Middle: Single-trial voltage traces (colored lines, 9-10 trials) and average response (black lines) to the first 10 s of the movie. Right: Output functions of LN model (40 bins) obtained from the responses to the natural movie. B. Scatter plot of the output nonlinearity index (ONi) measured under spatiotemporal white noise versus measured under the natural movies. Each black dot represents the average ONi over multiple movies of one bipolar cell (n=9), and gray dots show the ONi of individual movies (n=19). Black line shows the linear regression (**p<0.01), and dashed red line the identity line. C. LN model performance under natural movies for the same two sample cells as in A. Left: Trial-averaged response traces to the patterned spot (as in Figure 4). Right: Averaged response traces (black) to the first 10 s of the movie and the corresponding predicted (red) response traces from the spatiotemporal white noise LN model. The model performance (R^2^) is reported for the full 40 seconds of the movie. Black arrows for Cell 2 mark events that were not accurately predicted by the model. D. Population analysis of model performance. Top: Distribution of the model performance (R^2^) over all natural movies (n=19). Bottom: Scatter plot of R^2^ versus spatial nonlinearity index. Black dots represent the average R^2^ over multiple movies per cell (n=9, linear regression line in black, *p<0.05), gray dots the R^2^ for individual movies (n=19).

We first studied the contrast representation under natural statistics via the LN model. We used the spatiotemporal filter obtained from the white noise experiments and re-computing the output function from the responses to the natural movies (Heitman et al., 2016). The obtained output functions showed linear as well as nonlinear representation of contrast under natural movies in different cells (see sample cells in Figure 5A). We again quantified the degree of nonlinearity by computing the output nonlinearity index (ONi) as described before for white noise stimuli. The ONi under natural stimuli was strongly correlated to the ONi under white noise (Figure 5B; r=0.8, p=0.009, n=9 cells), and the two measures did not differ significantly (p=0.43, n=9 cells). Thus, the nonlinear contrast representation of bipolar cells remained similar under artificial and natural stimulus statistics.

To study spatial integration with natural movies, we investigated the performance of the LN model when predicting responses to these natural stimuli. Model predictions (Figure 5C) were obtained with the LN model as retrieved from spatiotemporal white noise (filter as well as output nonlinearity). Overall, the distribution of the model performance for natural movies (Figure 5D, top panel) had a similar spread as for the spatiotemporal white noise, ranging between 45-91% explained variance (n=19 movies; 9 cells). To test whether – as for the spatiotemporal white noise – it is the nonlinear spatial integration that is causing low prediction performance for some cells, we again studied the relationship between the spatial nonlinearity index (SNi) and the average model performance per cell. We observed a clear negative correlation (Figure 5D bottom panel, r=-0.73, p=0.027, n=9 cells). Thus, nonlinear spatial stimulus integration also limited the predictive quality of the LN model for some bipolar cells under natural stimuli.

### Input nonlinearities dominate the nonlinear characteristics of bipolar cells responses

Given that bipolar cells showed both nonlinear contrast representation (Figure 2) as well as nonlinear spatial integration (Figure 3), we asked whether these two nonlinearities are related. For example, the first sample cell in Figure 6A had an approximately linear representation of contrast under white noise stimulation as well as vanishing responses to the patterned spots, indicating also linear spatial integration. The second sample cell, on the other hand, displayed strongly nonlinear characteristics in both cases. Population analysis indeed revealed a positive correlation between the output nonlinearity index and the spatial nonlinearity index (r=0.57, p=0.017, n=17, Figure 6B); the higher the degree of nonlinearity in the contrast representation, the stronger the nonlinearity in the spatial integration.

**Figure 6.**
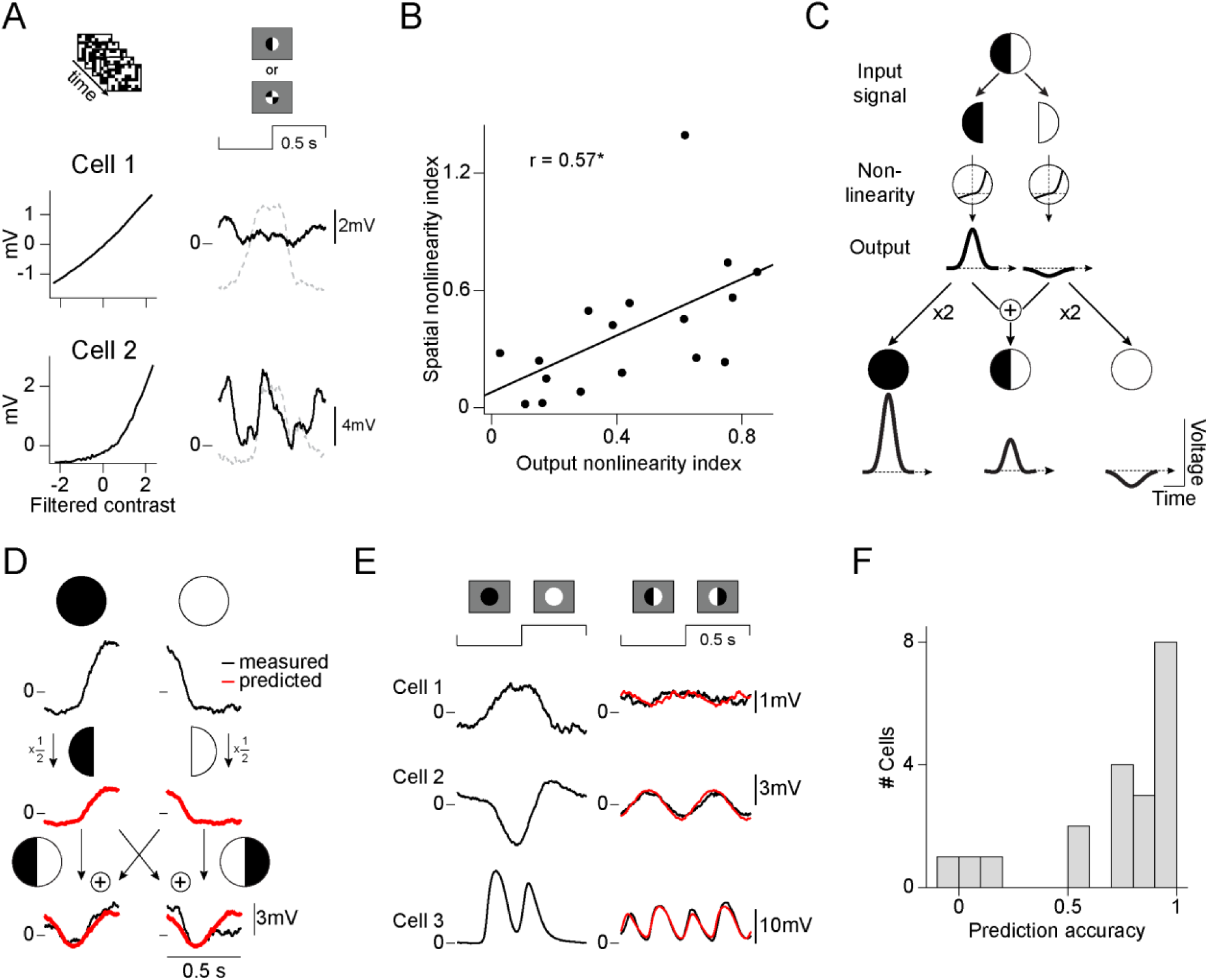
Input nonlinearities dominate the nonlinear characteristics of bipolar cells responses. A. Output functions of the white noise stimulus and trial-averaged response traces to the patterned spot (as in Figure 4) for two sample cells. B. Absolute value of the output nonlinearity index plotted against the spatial nonlinearity index. Each data point represents one bipolar cell (n=17, linear regression line in black, *p<0.05). C. Schematic view of the relationship between the responses to the split spot and the uniform spot for a model with only local nonlinearities. The input signals (half-field white and black stimuli) are passed through a nonlinear function to generate the corresponding output. For the split spot response (middle) the two outputs are summed; for a uniform spot, the corresponding single output is multiplied by two (left and right). D. Schematic depiction of the approach for predicting the response to the split spot with data from a sample cell. Top: Measured response to contrast-reversing uniform spots (black traces), split into two half-periods. Middle: Estimated response (red traces) to half-field black and white spots (obtained by multiplying the uniform-spot trace by 1/2). Bottom: Predicted (red) and measured response (black) to the contrast-reversing split spot. The response prediction is obtained by summing the responses to the half-field white and black stimuli for both split spot reversals. E. Trial-averaged response traces to uniform spots and split spots (black) together with the prediction for the split spots (red) for three further sample cells. F. Distribution of the prediction accuracy for the response traces to patterned spots (n=20).

The correspondence between the two measures of nonlinearity let us hypothesize that nonlinear signal transformations occur mainly locally, before spatial integration, and that the output nonlinearity is simply inherited from this transformation before the integration. If this is the case, then the responses to the split spot and to the uniform spot should be related through a simple model (Figure 6C). In this model, the contrast signal from each half of the receptive field is filtered and transformed by a local nonlinear function, and the two resulting signals are linearly summed to yield the membrane potential without any further nonlinearity. Therefore, in this model, the split spot response is the sum of the responses to a half-field white stimulus and a half-field black stimulus, which, in turn, are half of the responses to the full white and full black spots, respectively. This lets us predict the responses to the contrast-reversing split spot without having to explicitly take nonlinear transformations into account because the measured responses to the uniform spots already contain the local nonlinearities that affect the stimulus integration for the split spot. Note that the prediction for the patterned spot with four quarters is identical to that for the split spot because it effectively also comprises two receptive field halves of increasing and decreasing light intensity, respectively.

Figure 6D illustrates the procedure for the prediction of a sample cell that displayed nonlinear spatial integration. Indeed, we found that the predicted trace (red line) matched the measured response trace (black line) quite accurately. Figure 6E shows the responses and predicted traces of three further cells. For the first, the predicted response was small, similar to the cell’s actual response, which revealed linear spatial integration. The other two cells had larger response predictions, indicative of nonlinear spatial integration, matching the measured responses (see Figure S3 for more sample cells.) To quantify for all recorded cells whether this simple model captured the degree of nonlinearity revealed in the patterned-spot measurements, we computed a prediction accuracy measure, which assesses the deviation between predicted and measured response and takes a value of unity if the two match and zero for a mismatch (see Methods). For most of our cells (15 out of 20) we observed high prediction accuracy measures with values larger than 0.7 (see Figure 6F). These results suggest that the primary nonlinear signal transformations indeed happen before the integration of inputs by the bipolar cell and not at the level of the integrated bipolar cell membrane potential.

### Spatial scale of nonlinear signal processing in bipolar cells

In order to explore the mechanisms underlying the observed nonlinear transformations, we first investigated the spatial scale at which nonlinear integration appears. To do so, we analyzed bipolar cell responses to spot stimulation of receptive field centers with patterns of increasingly finer spatial structure. We used the split spots with two halves of opposing contrast as well as other patterned spots with divisions into quarters or into checkerboards with squares of 25 µm or 10 µm length (Figure 7A). As before, all patterns were contrast-reversed at 1 Hz. We found that fine checkerboards of 25 µm or smaller generally yielded no or very little responses (see Figure 7B), indicating that such fine spatial patterns are integrated linearly. By contrast, for subdivisions of the receptive field into quarters, responses were still nearly as strong as for the split spot, indicating that nonlinear spatial integration occurs at a scale of several tens of micrometers.

**Figure 7.**
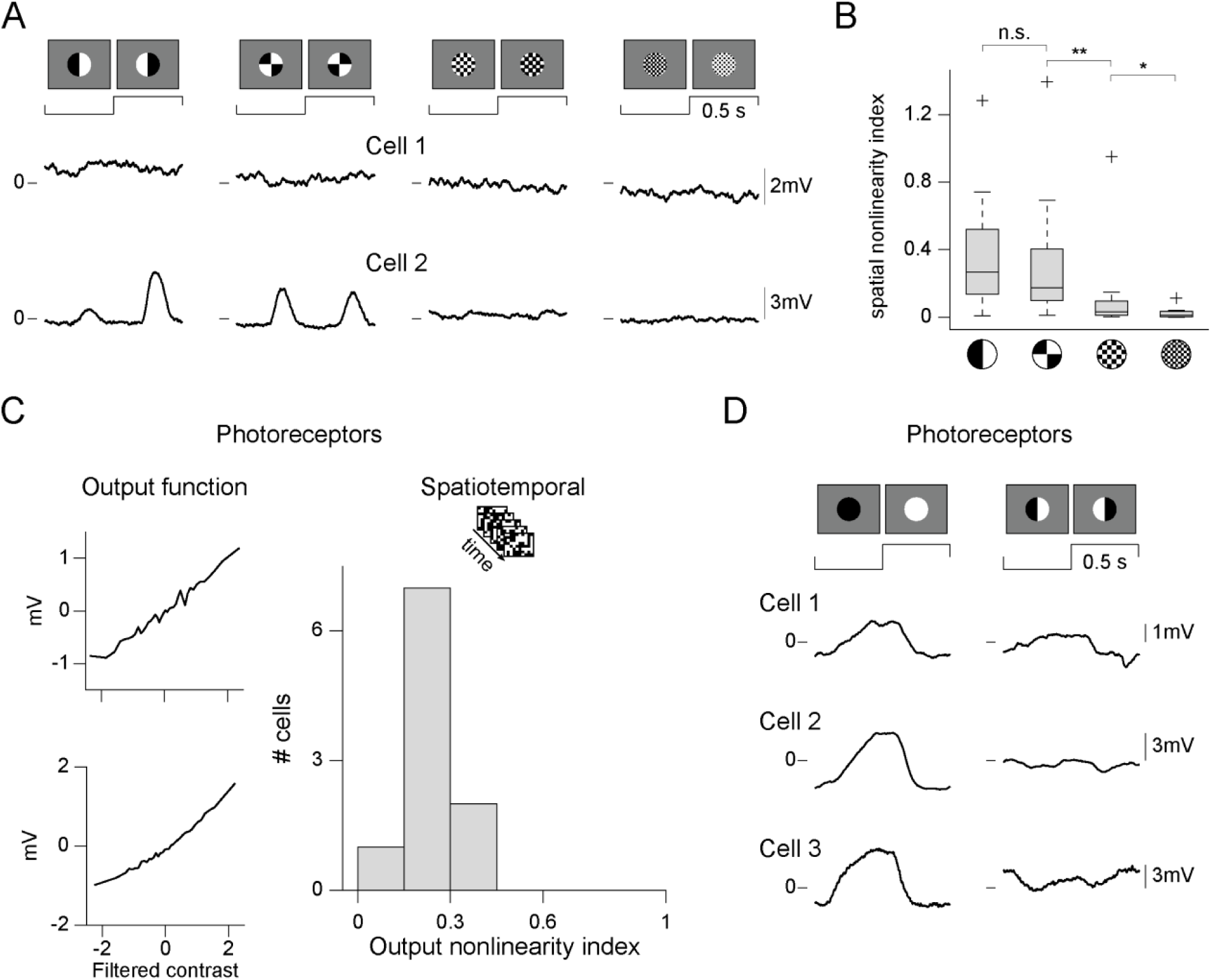
Spatial scale of nonlinear signal processing in bipolar cells. A. Trial-averaged response traces (as in Figure 4) of two sample bipolar cells to patterned spots of increasingly finer spatial structure. B. Distributions of the spatial nonlinearity index (SNi) for the different spatial structures: two halves (n=20), four quarters (n=20), 25 µm squares (n=20), and 10 µm squares (n=9). For each box, the central line marks the median and the edges the 25th and 75th percentiles, the whiskers mark the range of the data points and the crosses the outliers, **p<0.01, *p<0.05). C. Two sample photoreceptor output functions measured with spatiotemporal white noise and the distribution of the output nonlinearity index (ONi) for photoreceptors (n=10). D. Three recorded photoreceptor responses to the contrast-reversing uniform and split spots.

Thus, receptive fields might be organized into functional subunits of tens of micrometers in size, making it questionable whether photoreceptors with typical sizes of 10-15 µm in salamander (Mariani, 1986; Sherry et al., 1998) could be the source of the nonlinearity. Nonetheless, to directly test whether photoreceptors themselves display nonlinear stimulus encoding, we intracellularly measured their voltage responses in a similar fashion as for bipolar cells. We indeed found that photoreceptors, as compared to bipolar cells (cf. Figures 2D and 3B), showed more linear and less diverse output functions (Figure 7C) and also integrated signals linearly (Figure 7D).

### Relation of nonlinear processing and standard response properties in bipolar cells

It has been hypothesized that intrinsic properties of bipolar cells, such as the type of glutamate receptors at their dendritic tips (e.g., metabotropic versus ionotropic and AMPA versus kainate receptors), could contribute to nonlinear transformations (Demb et al., 2001). The receptor type furthermore determines the bipolar cell’s preferred contrast (ON versus OFF) and can also influence whether its light responses have transient or sustained characteristics (DeVries, 2000; DeVries and Schwartz, 1999; Euler et al., 2014; Wassle, 2004). We thus aimed at comparing these standard properties with the degree of nonlinearity on a cell-by-cell basis.

To assess whether a bipolar cell had transient or sustained response characteristics, we analyzed responses to spots of preferred contrast (see sample cells in Figure 8A) and quantified the response kinetics with a sustained-transient index (STi; see Methods), which takes values close to zero for transient and values close to unity for sustained responses. The STi was negatively correlated with the spatial nonlinearity index (Figure 8B, r=-0.76, p=0.0007, n=16), showing that more transient cells had stronger nonlinearities as revealed by the patterned-spot experiments. In line with the documented stratification pattern of transient bipolar cells (Awatramani and Slaughter, 2000; Baden et al., 2013a; Borghuis et al., 2013), we observed that cells with stronger spatial nonlinearities (i.e., higher spatial integration index) stratified more in the middle of the inner plexiform layer compared to cells with more linear spatial integration (see Figure S4A).

**Figure 8.**
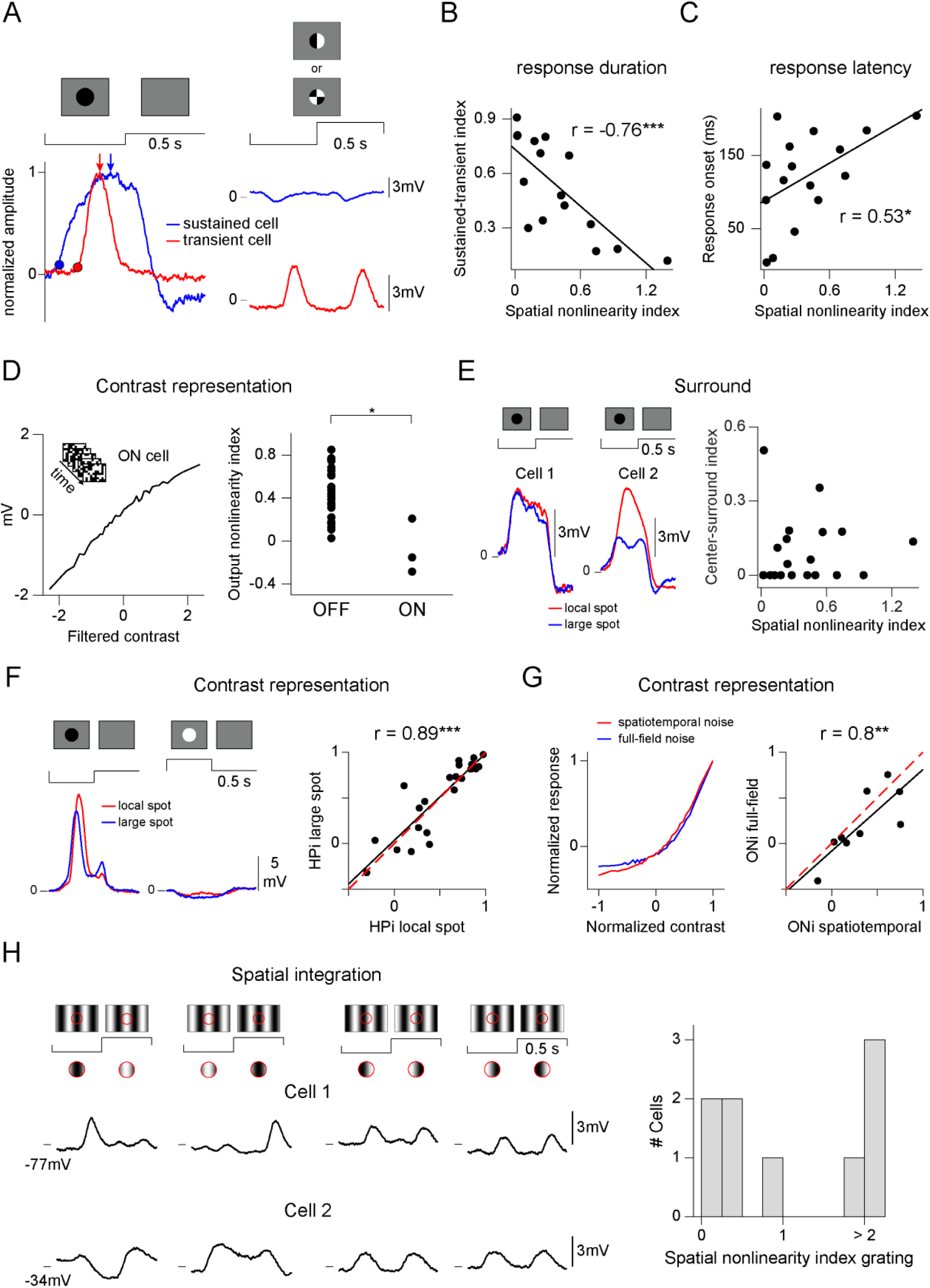
Relation of nonlinear processing and standard response properties in bipolar cells. A. Left: Trial-averaged response traces of two bipolar cells to uniform spots, normalized to the peak. The dots mark the response onsets and the arrows the response peaks. Right: Average response traces to the patterned spot for the same two cells (displayed as in Figure 4). B. Scatter plot of the spatial nonlinearity index (SNi) versus the sustained-transient index (STi). Each data point represents one bipolar cell. Black line is the linear regression line (n=16). C. Same as B for response onset. D. Example output function under spatiotemporal white noise of an ON bipolar cell that exhibits saturation (left) and comparison between the output nonlinearity index (ONi) for OFF and ON cells (right). Each dot represents one bipolar cell (26 OFF and 3 ON cells). E. Left: Trial-averaged response traces to local spots (red) and large spots (blue) for two sample cells. Right: Same as B for the center-surround index (n=20). F. Left: Trial-averaged response for a local (red) and large (blue) spot under preferred and non-preferred contrast for one sample cell (cf. Figure 2). Right: Scatter plot of the hyperpolarization index (HPi) for local versus large spots (n=23). The red dashed line shows the identity line. G. Left: Output function under spatiotemporal (red) and full-field (blue) white noise for one sample cell. The output functions were normalized to the maximum response, and contrast values (x-axis) were normalized to range from −1 to 1. Right: Scatter plot of the output nonlinearity index (ONi) for spatiotemporal versus full-field white noise (n=9). H. Left: Trial-averaged response traces of two bipolar cells to 4 phases of full-field reversing sinusoidal gratings. The red circle schematizes the stimulus pattern inside the receptive field. Right: Distribution of the spatial nonlinearity index under grating stimulation (n=9). (*p<0.05, **p<0.01, ***p<0.001)

We also checked whether nonlinear response characteristics were related to response latency. Latency can be measured as response onset or time-to-peak (Franke et al., 2017; Krieger et al., 2017), and we observed that sustained cells showed a fast response onset but longer time-to–peak, whereas transient cells showed a more sluggish onset but earlier peak (see example cells in Figure 8A and Figure S4B). Indeed, the two latency measures showed different correlations with the degree of spatial nonlinearity. Response onset was positively correlated with the spatial nonlinearity index (Figure 8C, r=0.53, p=0.03, n=16), whereas time-to-peak indicated a rather negative, yet non-significant correlation (r=-0.24, p=0.37, n=16). Thus, nonlinear bipolar cells are typically transient cells with a slow onset but perhaps faster peak response.

We also investigated whether there were systematic differences in nonlinear properties between ON and OFF bipolar cells. Note, however, that the salamander retina is biased towards OFF cells (Burkhardt et al., 1998; Hare et al., 1986; Hare and Owen, 1990; Segev et al., 2006), and our recordings yielded around 90% OFF and only 10% ON bipolar cells. The few ON cells showed comparatively smaller output nonlinearity indices with higher hyperpolarization than depolarization, which is reflected in a saturation of the output function, with negative output nonlinearity index values (p=0.016, n=29, Figure 8D).

Further, we checked whether the strength of the receptive field surround of a bipolar cell was related to the cell’s nonlinearity. We quantified the strength of the suppressive surround from the peak responses to large and small spots (Figure 8E) and used the response ratio as a center-surround index, which takes values close to zero for cells without surround and larger values otherwise (see Methods). When comparing the center-surround index to the spatial nonlinearity index, however, we did not find any correlation (r=0, p=0.95, n=20, Figure 8E); cells both with and without suppressive surround could show nonlinear properties. Consistent with this finding, the nonlinear nature of bipolar cells is not altered by how strongly the receptive field surround is stimulated. The relative amplitudes of hyperpolarization and depolarization for black and white spots were similar for small and large spots (r=0.89, p=2×10^−8^, n=23; Figure 8F), and the nonlinearity of contrast representation captured via the LN model (r=0.8, p=0.01, n=9, Figure 8G) did not depend on whether the stimulus was spatiotemporal or spatially homogeneous white noise, which activates the surround more strongly. In addition, we also tested full-field stimulation with classical reversing sinusoidal gratings (Enroth-Cugell and Robson, 1966) and found that they yielded similar results (in particular frequency doubling) as the local stimulation with reversing patterned spots (Figure 8H). Like the strength of the receptive field surround, several other investigated bipolar cell properties, such as the receptive field diameter, the occurrence of oscillations in the responses, or the shape of the temporal filter, also appeared to be unrelated to the nonlinear properties (Figure S4C-E).

## Discussion

We investigated the hypothesis of linear somatic light responses in salamander bipolar cells by intracellularly recording their membrane potential. We found nonlinear representation of contrast (Figure 2) as well as nonlinear spatial integration (Figure 3) in a sizable fraction of bipolar cells. Furthermore, under artificial and natural stimuli, linear receptive field models did not accurately describe the responses of spatially nonlinear bipolar cells (Figures 4 and 5). Our data suggest that nonlinear signal transformations arise before the input integration by the bipolar cell and that nonlinear representation of contrast by the membrane potential is largely inherited by these early transformations (Figure 6). Such nonlinear transformations might occur at the bipolar cell dendrites (transient vs. sustained) and most likely not in the photoreceptor voltage signals or in mechanisms of the receptive field surround (Figure 7 and 8).

### Nonlinear output of bipolar cells

Classically, bipolar cells had been assumed to produce a linear output because of the absence of action potentials in the membrane potential and because of the ribbon synapses at the bipolar cell synaptic terminals (reviewed by Shapley, 2009). Yet, later studies indicated nonlinear bipolar cell signals underlie nonlinear spatial integration in Y-type ganglion cells (Demb et al., 2001), and more recent observations directly showed nonlinear glutamate release as a function of contrast, especially at OFF bipolar cell terminals (Borghuis et al., 2013; Franke et al., 2017). Thus, bipolar cell signals are currently often pictured as being linear at the soma (i.e. equal gain for depolarization and hyperpolarization) and nonlinear, rectifying at the axon terminals with stronger increases in glutamate release under depolarization as compared to decreases in glutamate release for comparable hyperpolarization (Baccus et al., 2008; Borghuis et al., 2013; Roska and Meister, 2014). The view of linear somatic bipolar cell signals seems to be partly drawn from a few examples with measured linear responses under white noise or sinusoidal stimulation, mainly in the context of contrast adaptation (Baccus and Meister, 2002; Rieke, 2001; Sakai and Naka, 1987a, b; Toyoda, 1974). This appears to be inconsistent with our observations of nonlinearities in the contrast representation of artificial and natural stimuli (Figure 2 and Figure 5), though perhaps the limited number of cells in these previous reports together with the observed diversity of linear and nonlinear effects in different bipolar cells (Figure 8) may explain this seeming contradiction.

Several other studies in the salamander and mammalian retina had documented somatic nonlinearities, but often based on a much larger applied luminance range (e.g., scotopic to photopic) as compared to the present study (Burkhardt et al., 2011; Burkhardt and Fahey, 1998; Euler and Masland, 2000; Fahey and Burkhardt, 2003; Molnar et al., 2009; Wu et al., 2000). Under such large light intensity changes, nonlinear output functions are expected because of saturation in the changes of the membrane potential with changing light intensity. In the present work, nonlinearities were measured with smaller light intensity changes (e.g., as occurring within a single natural scene), indicating that nonlinearities are relevant not only when changing luminance regimes, but also for actual stimulus encoding within a natural range of Weber contrast values. Taken together, our results support a bipolar cell model where the voltage signals can be nonlinearly related to contrast and where additional rectification of the voltage signal occurs at the axon terminals (Demb et al., 2001).

### Nonlinear inputs to bipolar cells

Similar to the contrast representation, the signal integration in bipolar cells is viewed as linear because of linear photoreceptor responses and of the ribbon synapses at the photoreceptor terminals (Shapley, 2009). Support for linear signal integration comes from a small number of sample recordings in salamander bipolar cells that documented accurate response predictions to jittering gratings with a linear-receptive-field model (Baccus et al., 2008) as well as from observed response nulling in the glutamate release of mouse bipolar cell terminals under large reversing-gratings (Borghuis et al., 2013). Note, though, that the latter study focused on bipolar cells connected to alpha retinal ganglion cells whose major inputs likely come from few specific bipolar cell types (Schwartz et al., 2012; Tien et al., 2017; Yu et al., 2018), leaving open the question whether other bipolar cells in the mouse retina may differ in this respect. On the other hand, species-specific features might influence the observed nonlinearities. For example, the salamander retina has more photoreceptor types (Sherry et al., 1998) and the bipolar cells morphology might be more diverse (∼12-20 types; Pang et al., 2004; Wu et al., 2000) compared to the mouse retina (Euler et al., 2014).

It is worth noting that there is some evidence that nonlinear signal integration might exist in mammalian bipolar cells as well. For example, nonlinear signal integration has been found in rod bipolar cells in the mouse retina near absolute darkness (Berntson et al., 2004; Field and Rieke, 2002; Sampath and Rieke, 2004), though this is typically considered a rather specific scenario where the nonlinearity depends on the saturated state of the photoreceptor-to-bipolar cell signal transmission in complete darkness. More related to our investigations under photopic conditions, Freeman et al. (2015) observed that responses of primate ganglion cells did not completely cancel out when two cones that projected onto the same bipolar cell were stimulated with opposing contrast. They attributed the remaining response to nonlinearities in cone photoreceptors, which would lead to nonlinear integration by the bipolar cell, in line with our findings. Similarly, some ganglion cells in mouse retina are sensitive to patterns with spatial scales of around 20 µm (Jacoby and Schwartz, 2017; Krieger et al., 2017; Mani and Schwartz, 2017; Schwartz et al., 2012; Zhang et al., 2012), though this high spatial sensitivity was interpreted as resulting from spatially linear bipolar cells with small receptive fields and nonlinear synaptic release. Yet, given that reports of receptive field sizes of mouse bipolar cells range from 40-80 µm (Berntson and Taylor, 2000; Borghuis et al., 2013; Franke et al., 2017; Schwartz et al., 2012), spatially nonlinear bipolar cells could provide a viable alternative source of the observed spatial sensitivity.

### Putative mechanisms for nonlinearities in bipolar cells

The observed nonlinearities appear to arise mainly from mechanisms occurring before the signal integration by the bipolar cell in the soma (Figure 6). Such nonlinear signal transformations could occur in dendritic branches of the bipolar cells (Demb et al., 2001). There, receptors diversify the photoreceptor signals into ON and OFF (metabotropic vs. ionotropic receptors) and transient and sustained (AMPA vs. kainite receptors) bipolar cell channels (DeVries, 2000; DeVries and Schwartz, 1999). Along this line, we found that transient bipolar cells showed more nonlinear integration properties compared to sustained bipolar cells (Figure 8B) and only ON bipolar cells showed response saturation (Figure 8D). Thus, differences in the receptor dynamics or dendritic ion channels – e.g. voltage-dependent sodium channels (Zenisek et al., 2001) – of the different cell types could lead to the observed nonlinear input signals.

Another potential source of nonlinear signals could be the photoreceptors’ synaptic terminals. Nonlinear calcium signals have indeed been observed at some mouse cone terminals (Baden et al., 2013b), which may still be consistent with linear somatic voltage signals of photoreceptors as found here (Figure 7C-D). However, the observed linear output functions in photoreceptor were less diverse compared to the bipolar cells (compare Figure 2D and 7C). And the fact that different bipolar cells could display quite different levels of nonlinear integration appears incompatible with a generic mechanism in photoreceptors, which should affect all bipolar cells similarly, indicating that the nonlinear signal transformation likely happens after the photoreceptors.

The nonlinear signal transformations could, in principle, also occur from interactions with other cells. For example, amacrine cells act on bipolar cells terminals. We minimized the influence of amacrine cells by recording the voltage of bipolar cells at the soma (before the terminals). Yet, bipolar cells are compact neurons and the signals at the terminals might back-propagate to the soma and influence the somatic recordings (Euler and Masland, 2000; Masland, 2012). However, amacrine cell feedback modulates glutamate release from bipolar cells mainly for stimuli considerably larger than the bipolar cell receptive field center and may have little effect for local stimuli (Franke et al., 2017), whereas we observed similar nonlinear properties under small and large stimuli (see Figure 8E-H). Furthermore, our observed nonlinear input integration in bipolar cell manifested itself as depolarization events, which in addition originate before signal integration in the soma, thus making a mechanism through inhibitory input at the terminals unlikely.

For similar reasons, also horizontal cells and the rarely observed interplexiform cells, which are located in the inner retina and provide glycinergic input onto the dendrites of bipolar cells (Maple and Wu, 1998), are unlikely to generate the nonlinearity. Activating glycine receptors through interplexiform cells hyperpolarizes OFF bipolar cells (Jiang et al., 2014), whereas the nonlinear responses to patterned-spot stimulation consisted of depolarization events. And horizontal cells have large receptive fields (500 to 1600 µm) in the salamander retina (Lasansky and Vallerga, 1975; Zhang et al., 2006), and thus, they are expected to be more activated under global compared to a local stimulation (Figure 8E-H).

### Implications of nonlinear bipolar cells for computational models in the retina

For retinal ganglion cells, linear receptive field models often fail to describe the responses to artificial and natural stimuli (Freeman et al., 2015; Heitman et al., 2016; Liu et al., 2017; Schwartz et al., 2012; Turner and Rieke, 2016). Here, we observed such a failure already for bipolar cells, upstream of ganglion cells (Figures 4 and 5). Spatially nonlinear models (LNLN models) might improve the response prediction of bipolar cells. The observed connection between the contrast representation (output nonlinearity) and the spatial integration (Figure 6) could help in constraining the model structure and parameters.

For retinal ganglion cells, several studies already investigated spatially nonlinear models (so-called subunit models or LNLN models) by subdividing the receptive field into multiple subfields (subunits at the size of bipolar cells receptive field) and transforming the inputs in each subunit nonlinearly (e.g. rectification) before summation (Freeman et al., 2015; Liu et al., 2017; Schwartz et al., 2012; Turner and Rieke, 2016). It may be an interesting endeavor to see whether adding another layer of nonlinear integration, reflecting the bipolar cell nonlinearities inside the subfields of the ganglion cell receptive field, could improve the often still unsatisfactory response predictions of LNLN models for retinal ganglion cells.

### Functional relevance of nonlinear computations in bipolar cells

Many retinal ganglion cell types are thought to extract diverse features from the visual scene. This feature extraction depends on subunits within the ganglion cell receptive field, which nonlinearly transform contrast information so that local signals do not cancel out (Baccus et al., 2008; Demb et al., 2001; Freeman et al., 2015; Gollisch, 2013; Gollisch and Meister, 2010; Hochstein and Shapley, 1976a; Liu et al., 2017; Schwartz and Rieke, 2011; Schwartz et al., 2012). Excitatory inputs have been documented to be the substrate for the nonlinear subunits (Demb et al., 2001), and consistent with those findings, we have observed nonlinear contrast representation in bipolar cells.

In addition, we have observed nonlinear spatial integration in bipolar cells, which could provide ganglion cells with feature sensitivity on a spatial scale below the size of bipolar cell receptive fields (Freeman et al., 2015). In dim light conditions near complete darkness, for example, retinal ganglion cells as well as human observers can report the detection of single photons by rod photoreceptors (Barlow et al., 1971; Hecht et al., 1941). The high sensitivity comes through nonlinear signal integration in rod bipolar cells (Berntson et al., 2004; Field and Rieke, 2002; Sampath and Rieke, 2004). Our findings could point to a similar functional mechanism during daylight where local changes that happen below the scale of bipolar cell receptive fields are reported to ganglion cells and the brain. This maintains sensitivity to small spatial structures despite the signal convergence from photoreceptors to ganglion cells and could be particularly relevant for animals whose retinal cells have large receptive fields in terms of visual angle, e.g. because their eyes are small.

## Experimental procedures

### Electrophysiology

All experimental procedures were approved by the University Medical Center Göttingen. The eyes of dark-adapted (∼1h) axolotl salamanders (*Ambystoma mexicanum*; pigmented wild type, both sexes, n=20) were enucleated, and the vitreous humor was carefully removed. In 19 experiments, the whole retina was detached from the pigmented epithelium, mounted over a hole of 1.5-2 mm diameter on a nitrocellulose filter membrane and placed ganglion cell-side-down on a 60-channel perforated multielectrode array (MEA) (Reinhard et al., 2014). A pump outside the setup applied slight suction through the small holes of the perforated MEA to keep the retina in place, which permits access to the cells from the photoreceptor-side for intracellular recordings (see Figure 1A). In one experiment, the eyes were hemisected and the retinas were placed ganglion cell-side-down into the bath chamber on a nitrocellulose filter membrane without a MEA. During the experiment, the retina was superfused with oxygenated (95%O_2_ and 5%CO_2_) Ringer’s medium (110mM NaCl, 2.5mM KCl, 1mM CaCl_2_, 1.6mM MgCl_2_, 22mM NaHCO_3_, 10mM D-Glucose monohydrate) at ∼25°C.

To obtain intracellular recordings from bipolar cells, we used sharp glass microelectrodes (BF120-60-10; shaped with a P-97 Brown/Fleming pipette puller, Sutter Instruments), which allowed penetrating the retina with little damage and provided for fairly long recordings (∼0.5-2 hours). Such long recordings were attainable in the salamander retina because of the comparatively large cell bodies. Microelectrodes were tip-filled with 4% Neurobiotin (dissolved in 0.1M Tris buffer) and backfilled with 3 M KCl solution (resistance 206 ±85 MΩ, mean ±SD). With the help of a 60x objective, the tip of the microelectrode was placed above the outer segments of the photoreceptors over a MEA recording site that showed high spiking activity to maximize chances of observing ganglion cell responses under current injection into the bipolar cell. Inside the bath solution, the pipette offset was nulled and the microelectrode was slowly inserted into the retina (1-µm steps coaxially to the electrode) until a cell was impaled. The depth of the electrode tip was monitored with the remote control of the micromanipulator (keypad SM5, Luigs & Neumann). The membrane potential of the cell was recorded using a MultiClamp 700B amplifier (Molecular Devices, San Jose, CA) and digitized at 20 kHz (Digidata 1440A, Molecular Devices). For all the described data analyses, the recorded voltage traces were further filtered by a running median (window=80 data points) and downsampled to 1-ms resolution by taking every 20th data point.

### Cell-type identification

To assess the morphology of the intracellularly recorded cells, Neurobiotin was injected at the end of a recording with current pulses (blocks of positive and negative pulses of 80-200 pA, 0.5 s current and 2 s break) for 3-7 minutes. Afterwards, the retina was carefully removed from the array, fixated with 4% formaldehyde and further processed with Alexa Fluor 488 Streptavidin (Thermo Fisher Scientific, MA, USA) and with To-Pro-3 (Thermo Fisher Scientific, MA, USA) for nucleus staining. The Neurobiotin-filled cells were imaged with a confocal microscope, and two-dimensional representations were constructed with maximum projections over manually chosen regions to best represent the dendrites and axon terminals within the inner and outer plexiform layer (horizontal views, x-y planes) and the vertical view across the bipolar cell (representation in x-z or y-z planes). For Figure S4A, we traced the cell with the semi-automatic Fiji plugin Simple Neurite Tracer and marked the boundaries of the inner plexiform layer by detecting the nuclei of the ganglion cell layer and of the inner nuclear layer (INL) with the help of To-Pro-3 nucleus staining. The traced morphologies were further analyzed with Matlab, and the axon stratification depth within the inner plexiform layer (IPL) was computed as the distribution of the relative distances along the axon branches to the INL boundary, normalized so that the boundary to the ganglion cell layer is at a distance of unity (Wu et al., 2000).

When the staining and imaging were successful, bipolar cells were identified based on their bipolar shape, with neurites in the outer and inner plexiform layer (see Figure 1B). Photoreceptors, on the other hand, could be well recognized by their outer segments, and amacrine cells showed neurites only in the inner plexiform layer (see Figure S1). In some experiments, the staining failed, or we could not remove the entire retina from the perforated MEA. Thus, we developed additional criteria to identify bipolar cells. We distinguished bipolar cells (BC) from photoreceptors (PR) by the recording depth (morphologically identified PR were observed directly when entering into the retina, mainly at a depth <100µm, BC at a depth of 135±40 µm, mean ± SD), the receptive field size (morphologically identified PRs showed smaller RFs, 34±17 µm, than BCs, 76 ±27 µm, mean ± SD), and the characteristic visual response of photoreceptors to contrast steps. Bipolar cells were distinguished from amacrine cells by observing the response polarity of the simultaneously recorded ganglion cells to positive and negative current pulses (50-500 pA, 500 ms duration, 2 s interval, for 2-4 minutes), which were injected into the intracellularly recorded cell. The spikes of the ganglion cells were extracted by a semi-automatic custom-made spike sorting program, based on a Gaussian mixture model and an expectation-maximization algorithm (Pouzat et al., 2002). If the majority of the retinal ganglion cells responded repeatedly to positive current injection, the intracellular recorded cells were identified as bipolar cells (Asari and Meister, 2012, 2014). If, however, the majority of the ganglion cells responded to negative current, the intracellular recorded cells were identified as amacrine cells (Asari and Meister, 2014; de Vries et al., 2011). In total, we identified 35 cells as bipolar cells and focused our analysis on 32 cells that showed clear preference to either negative (OFF cells) or positive (ON cells) contrast steps. The remaining 3 cells were excluded due to unclear contrast preference.

### Visual stimulation

Visual stimuli were generated by a custom-made software written in C++ and OpenGL and presented on a monochromatic gamma-corrected white OLED monitor (eMagin, 800×600 pixels, 60 or 75 Hz refresh rate). The image of the OLED screen was combined with the light path of an upright microscope through a beamsplitter and focused through a custom-made optics system and the 4x objective of the microscope onto the photoreceptor layer. The pixel resolution at the photoreceptor layer was 2.5 µm x 2.5 µm, and the mean light intensity was 2.5 mW/m^2^ in the low photopic range.

### Analysis of responses to spot stimuli

During the experiment, we first estimated the location over which subsequent spot stimuli were presented (online-determined center parameters). To do so, we shifted a contrast-reversing spot, whose diameter could be manually increased or decreased, over the screen, until a position and diameter size were found that maximally stimulated the cell (n=17). Alternatively, we applied online analysis of responses to spatiotemporal white noise to obtain the receptive field parameters (n=6; see below).

#### Estimation of optimal spot size (“local spot”)

To estimate the spot size that best stimulated a cell’s receptive field center (“local spot”) for further analyses, we presented spots of black and white contrast steps (100% contrast) at 11 (n=16) or 13 (n=3) fixed diameters (between 10-1200 µm) in random order, centered over the online-determined location. The spots were presented for 0.5 s on a background of mean light intensity and separated by 1 s of background illumination. Each spot size was presented on average 8 times. For each trial, we subtracted the average membrane potential measured over the 200 ms prior to the spot, and for each spot size, we averaged the baseline-subtracted responses over trials. For 4 cells, we presented the spots without an interval of background illumination between individual spots and only 6 spot diameters (50, 100, 200, 300, 400, 500 µm). These spots contrast-reversed (100% contrast) at 1 Hz for 4 s, with different spot sizes separated by background illumination of 4 s, which was used for computing the baseline membrane potential.

We used the peak of the trial-averaged response per spot size of the preferred contrast and fitted a difference-of-Gaussians model as described by Borghuis et al. (2013):

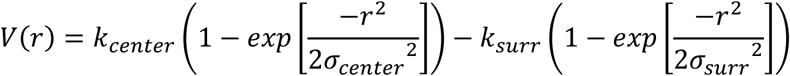

Here V is the peak membrane potential, r is the spot radius, k_center_ and k_surr_ are the maximum response amplitudes of the center and surround, and σ_center_ and σ_sur_ parameterize the radius of center and surround. For further spot-based analyses (see below), we then defined the local spot as the spot from the set of tested spot sizes with radius closest to an estimated receptive field width of 1.5 σ_center_ (diameter=2*1.5 σ_center_; 96 µm±36 µm, mean±SD, n=20). For 3 cells, we observed no response saturation or suppression for larger spot diameters but a rather continuous increase in response with every increase in spot size. For those 3 cells, the estimated receptive field width from the fit was much larger than for the other cells (>250 µm). Thus, to study the responses to a local spot stimulus, we chose for those 3 cells the closest spot size to the diameter computed with spatiotemporal white noise (see below), which was between 100-160 µm.

#### Hyperpolarization index

We analyzed the contrast representation with the local spot determined as described above (Figure 2) as well as with a large spot (500 µm diameter; Figure 8F). For both local and large spots, we computed a hyperpolarization index (HPi) by comparing the peak membrane potential during preferred contrast (depolarization), *V*_*dep*_, and the minimum membrane potential during non-preferred contrast (hyperpolarization), *V*_*hyp*_:

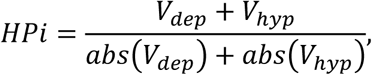

where *abs*(·) stands for taking the absolute values. The index takes values close to zero for cells that showed equal amounts of hyper- and depolarization, values near unity for cells that did not show any hyperpolarization (i.e. cells with rectified contrast signaling), and occasionally negative values for cells that showed stronger hyperpolarization than depolarization.

#### Sustained-transient index

We computed the sustained-transient index (STi) from the responses to the local spot of preferred contrast (described above). The STi was defined as the ratio of the steady-state response (average membrane potential over the last 50 ms of the spot presentation, *V*_*steady state*_) and the peak response (*V*_*peak*_):

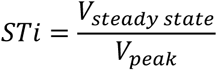

Cells with sustained responses (steady state≈peak) have an index near unity, whereas cells with transient responses (steady state≈0), have an index near zero.

#### Latency

We computed the latency from the local spot of preferred contrast (same as for the STi). We computed two measures of latency: the response onset and time-to-peak. The response onset (Franke et al., 2017) was defined as the time of the first data point after the spot onset that exceeded 3 standard deviations of the baseline (measured during the last 200 ms of the preceding background illumination). The time-to-peak was defined as the time from stimulus onset to the time of maximum response (Krieger et al., 2017).

#### Center-surround index

We computed the center-surround index from the spot stimuli described above by comparing the peak response over all spot sizes (up to including 500 µm, *V*_all spots_) with the peak response for a large spot of 500 µm (*V*_large spot_):

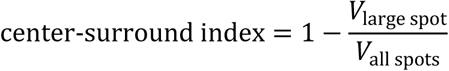

The center-surround index takes values close to zero if the response to the large spot was similar to the maximum response over all spots and values larger than zero if the response was reduced for the large spot.

#### Spatial nonlinearity index

To test for spatially nonlinear stimulus integration, we presented uniform and spatially structured spots (“patterned spots”) at the online-determined center parameters (see above). The patterned spots were obtained by dividing the uniform spot into two halves (“split spot”), four quarters, or into a checkerboard layout with squares of 25 µm or 10 µm, with opposite contrast (±100%) in adjacent stimulus subfields (see Figure 7). The uniform spot and the patterned spots were periodically reversed at 1 Hz for 4 seconds, followed by 4 seconds at background illumination. For each voltage trace, the baseline, determined as the mean voltage over the last 200 ms of the preceding background illumination, was subtracted.

To analyze the responses, we computed the average response for one stimulus cycle of 1 s duration (leaving out the first cycle to reduce stimulus onset artifacts), subtracted the mean, and performed a Fourier analysis (Matlab function *fft*) to obtain a power spectrum. If cells integrated the stimulus over space nonlinearly, both reversals of the spatial patterns could activate the cell, leading to frequency doubling (power at 2 Hz) in the response (Hochstein and Shapley, 1976b). For some cells, we further observed a 4-Hz component in the response to the patterned spots (see Figure S2B). We therefore computed a spatial nonlinearity index (SNi) for each spatial pattern from the combined power in relevant higher harmonics (2 Hz and 4 Hz) of the spatially structured spot, normalized by the power at 1 Hz of the uniform contrast-reversing spot:

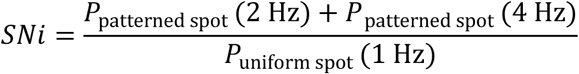

The normalization term acts as a generic measure of response strength of the cell (Turner and Rieke, 2016). Note that for 8 out of 20 cells, the uniform contrast-reversing spot used for normalization was presented with a fixed set of diameters (50, 100, 200, 300, 400, 500), which did not contain the exact diameter size of the patterned spots. We therefore linearly interpolated the membrane potential traces from the two uniform spots closest to the diameter size of the patterned spots and then computed the 1-Hz power to obtain the normalization term. Finally, we chose the maximum SNi over all spatial patterns as a representation of the cell’s integration property (Hochstein and Shapley, 1976b). A resulting SNi value near zero means that the cell did not show substantial frequency doubling for any spatial pattern, indicating linear spatial integration.

To study the integration properties under global stimulation, we presented full-field spatial sine-wave gratings with spatial periods of 80, 160, and 300 µm and 8 equally spaced spatial phases each. The gratings were reversed at 1 Hz for 8 seconds. For each spatial phase, we computed the average response per cycle (leaving out the first cycle to reduce stimulus onset artifact) and computed the Fourier transform as described above to assess the power at 1 Hz and higher harmonics (2 and 4 Hz). For each spatial period, we computed a spatial nonlinearity index (SNi_grating_) by summing the mean power at 2 Hz and 4 Hz, averaged over all phases, and dividing the sum by the maximum power at 1 Hz over all phases (Hochstein and Shapley, 1976b; Petrusca et al., 2007). We chose the maximum SNi_grating_ over all spatial periods to quantify how a cell integrated contrast under global stimulation (Hochstein and Shapley, 1976b).

#### Prediction of responses to the split spot

To predict the response to one reversal of the split spot experiment, we multiplied the averaged trace of the uniform contrast-reversing spot by 0.5 and summed the parts from the first 500 ms (black) and the second 500 ms (white). The prediction for a full cycle of the reversing split spot was then obtained by concatenating two identical 500-ms predictions for a single reversal. Note that the predicted trace was identical for the different spatial patterns (spot with two halves, four quarters, 25 µm or 10 µm checkerboard), because the total amount of black and white contrast inside the receptive field stayed the same. Note also that we did not need to adjust for response latency, which delays the response to the black or white spot compared to the time of the contrast reversal, because the same delay is expected for the reversing patterned spot. The response latency thus simply results in the same temporal phase shift of the periodical signals of the predicted and measured responses under reversing patterned spots.

For comparison with the predicted trace of a given cell, we selected the response trace of the patterned spot with the highest spatial nonlinearity index because this indicated the most balanced activation by both reversals. To quantify the similarity between the predicted response trace, *V*_predicted_(*t*), and the measured response, *V*_patterned spot_(*t*), of the cell, we first shifted the predicted response to have the same mean as the measured response in order to account for drifts in the baseline. Then we computed the prediction accuracy as follows:

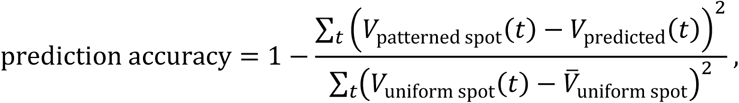

where *V*_uniform spot_(*t*) is the response trace for reversing uniform spots and 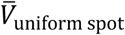 the corresponding temporal average. The measure is similar to the coefficient of determination, and the adjusted denominator serves to normalize the deviation between predicted and measured by the scale of a generic response rather than by the response to the patterned spot itself, which may be near zero for a linear cell. The similarity measure takes values near unity if the predicted and measured responses match and values near zero if they are unrelated.

### Spatiotemporal white noise analysis

#### Receptive field estimation

We visually stimulated the retina with binary spatiotemporal white noise in a checkerboard layout, where each square had a size of 30 µm and was updated randomly to black or white (100% contrast; denoted as stimulus values of ±1 for analysis) at 30 (n=8), 15 (n=8), 10 (n=2), or 7.5 Hz (n=1). For cells for which we also recorded natural movies, the squares had a size of 22.5 µm and were updated at 25 Hz (n=11) or 12.5 Hz (n=2) to fit the spatial and temporal resolution of the natural movies (see natural movie subsection below). Recording duration under spatiotemporal white noise was 23 ± 19 minutes (mean ± SD). To remove slow fluctuations in the responses, the voltage traces were first de-trended with a high-pass filter (Butterworth filter, 0.1 Hz cutoff) and then binned at the temporal resolution of the stimulus by computing the average membrane potential per time bin.

The spatiotemporal receptive field was computed in the following way: For each time bin, the average membrane potential was used as a weight for the preceding stimulus sequence over two seconds to compute a response-weighted average of all 2-s stimulus sequences, analogous to the common calculation of the spike-triggered average for spiking neurons (Chichilnisky, 2001). From the obtained response-weighted average, we determined the pixel with the largest absolute value over space and time, selected a window around the pixel of 720 µm to the side, and separated the response-weighted average within this window into the highest-ranked spatial and temporal components by singular-value decomposition (Gauthier et al., 2009).

To extract the receptive field location and size, we fitted a two-dimensional Gaussian function to the spatial component. We used the 1.5-sigma contour of the fitted Gaussian for displaying outlines of receptive fields and approximated the receptive field diameter as the diameter of a circle with the same area as within the 1.5-sigma contour. For some cells, we additionally recorded binary white noise of 10 µm x 10 µm square sizes to better separate the small receptive field contours of photoreceptors from bipolar cells. We excluded 3 cells out of 32 cells from the spatiotemporal white noise analysis because of noisy receptive fields. We detected noisy receptive fields by computing the average pixel intensity within the 3-sigma boundary of the Gaussian fit from the frame that contained the maximum pixel intensity and checking whether this signal was smaller than the noise level, determined as 3 standard deviations of the values in the spatiotemporal receptive field in the window 2-4 seconds before the spike.

#### Output nonlinearity index

To assess the degree of nonlinear contrast representation under white noise stimulation, we analyzed the output function (“nonlinearity”) of the linear-nonlinear (LN) model (see Figure 2). The first stage of the model is the linear spatiotemporal filter. To avoid noise contributions from pixels outside the receptive field, we reduced the number of elements in the spatiotemporal filter by setting pixel values of the response-weighted average outside the 3-sigma contour of the Gaussian fit to zero and re-separating the response-weighted average within this window into the highest-ranked spatial and temporal components by singular-value decomposition (as described above). Each component was normalized to unit Euclidean norm. We assumed space-time separability and applied the spatial and temporal component of the response-weighted average as separate filters. This yielded good approximations of the full spatiotemporal response-weighted average and helped avoid overfitting by strongly reducing the number of filter parameters. To obtain the output function, we first applied the spatial filter to each frame of the white noise stimulus by computing the scalar product between the spatial filter and the frame’s pixel contrast values and then convolved the resulting temporal sequence with the temporal filter to obtain the linear prediction of the LN model, also called the generator signal. Finally, the output function was obtained as a histogram by binning the generator signal values into 40 bins with equal numbers of data points and averaging the generator signal as well as the corresponding membrane potential values for each bin.

To quantify the degree of nonlinearity in the output function, we computed an output nonlinearity index (ONi) by fitting straight lines separately to the right half of the output function (positive generator signal values) and to the left half (negative generator signal values) and comparing the corresponding slope values *S*_pos_ and *S*_neg_:

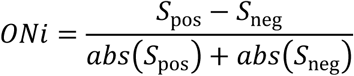

The index takes values close to zero when the two slopes were identical, indicating a linear representation of contrast, whereas values near unity correspond to nonlinear thresholding (i.e. rectification) of negative values of the generator signal. Negative values indicate a weaker response to positive generator signals, which can occur for saturating responses (see Figure 8D).

#### Assessing LN model performance

For 26 cells, we used a non-repeating binary white noise sequence, where 200 segments of 300 stimulus frames duration were randomly chosen as test data. For each test data segment, the LN model (filter and output function) was obtained from the remaining data (training data). For each stimulus frame of the test segment, the membrane potential was predicted by filtering the preceding stimulus sequence and applying linear inter- and extrapolation of the output function to extract the corresponding membrane potential value. The measured and predicted responses were compared by computing the explained variance (R^2^) as the square of the Pearson correlation coefficient R, and performance was reported as the average R^2^ over all test data segments. For the response traces in Figure 4B, the test segment is shown for which R^2^ was closest to the average R^2^.

For 6 cells, we used a non-repeating binary white noise sequence that was regularly interrupted (every 1200 frames) with an identical sequence of 300 frames (test segment, 10 trials on average). The LN model was obtained from the non-repeated training sequence, and we predicted the response to the test segment. The prediction performance was computed as R^2^ between the predicted and average measured response trace.

#### Temporal filter latency

The time-to-peak of the temporal filter was approximated by fitting a second-order polynomial to 3 data points (at stimulus bin resolution) around the maximum (for ON cells) or minimum (for OFF cells) of the filter and selecting the extremum of the fit (Khani and Gollisch, 2017).

#### Biphasicness index

Temporal filters were often biphasic with a second peak at longer latency and opposite sign as compared to the first peak. To quantify this, we calculated a biphasic index as the ratio between the absolute values of the second and first peak of the temporal filter (Khani and Gollisch, 2017; Zaghloul et al., 2007), where the first peak was defined as the maximum for ON cells and minimum for OFF cells and the second peak was the extremum of opposite sign. The index takes values close to zero when the second peak was close to zero (monophasic filter shape) and values of unity when the second peak had the same size as the first peak.

### Full-field white noise analysis

For 11 cells, we recorded responses to full-field white noise, which homogeneously activates the entire receptive field in a global fashion. Light intensity values were randomly drawn from a Gaussian distribution with a standard deviation of 30% around the same mean light intensity as for the other stimuli and updated at 30 Hz (n=9) or 25 Hz (n=2). The stimulus was composed of a non-repeated intensity sequence (training set), which was regularly interrupted (every 900 frames) by the same, repeated sequences of 300 frames (test set, presented on average 13 times). For analysis, light intensity values were converted to Weber contrast. The linear filter and output function of the LN model were obtained from the training set and computed in the same way as for the spatiotemporal white noise, except that the stimulus sequence was one-dimensional and the filter therefore only had a temporal component. The linear filter was normalized to unit Euclidean norm. In Figure 8G, we computed the output nonlinearity index from the output function of the full-field white noise in the same way as described above for spatiotemporal white noise. To assess the performance of the LN model, the response to the test set was predicted by first convolving the test stimulus sequence with the filter and then linearly inter- and extrapolating the output function to obtain the corresponding membrane potential. The performance of the LN model was again quantified by the explained variance (R^2^) between the predicted and the average measured response trace.

### Natural movie analysis

The natural movies were chosen from the “CatCam” database, where diverse outdoor scenes (e.g. woods, grass) had been recorded with a camera mounted on the head of a cat (Betsch et al., 2004) at 25 Hz and a resolution of 320×240 pixels. The movies had been used previously to study responses in ganglion cells, visual cortex, and lateral geniculate nucleus (Katz et al., 2016; Kayser et al., 2003; Mante et al., 2008). We chose five different movies of 20-40 seconds duration. For the projection on the retina, movie pixels spanned 3×3 pixels of the display projector to match the resolution of the spatiotemporal white noise, and movies were then cropped to the 800×600 pixels of the display. The light intensity was scaled so that the mean intensity was the same as for the other visual stimuli, and the standard deviation of pixel intensities was near 45% of the mean intensity. Movies were displayed at 25 Hz (except for two cells where the frame rate was reduced to 12.5 Hz) and repeated 9 times on average.

To predict the responses to the natural movies, we applied the LN model as obtained under binary white noise stimulation. The series of frames of the natural movie, represented by the contrast values relative to the overall mean intensity of the movie, was filtered with the spatial and temporal filters, and the filter output was passed through the corresponding nonlinearity, using inter- and extrapolation, as described above. The model performance was measured by the R^2^ between the averaged movie response and the prediction. From a total of 13 cells, we excluded 3 cells with noisy receptive fields (see spatiotemporal white noise section). Additionally, we excluded 1 cell and the responses to 2 movies from 2 cells, where the baseline membrane potential under the natural movie (average over the last 10 seconds) had increased by more than 20 mV compared to the starting voltage level under spatiotemporal white noise (average over the first 10 seconds). For some cells, we observed a drift in the overall response amplitude between the natural movie and the spatiotemporal white noise. For better visual comparison of the temporal profile of response and prediction, we therefore corrected for those offsets by normalizing the displayed predicted traces in Figure 5C to have the same mean and standard deviation as the response to the natural movie.

To quantify the nonlinearity of contrast representation under natural movies, we computed an output nonlinearity directly for the natural movies in the same way as for the white noise data, but using the temporal and spatial filters obtained from white noise (Heitman et al., 2016). Thus, we related the filter output when using the movie as a stimulus to the corresponding measured membrane potential via a histogram (40 bins). The degree of nonlinearity was then quantified by computing the ONi for this output function as explained above. Note that this nonlinearity obtained directly from the natural movie was not used for the response prediction via the LN model.

### Oscillation frequency

To analyze oscillations that were observed in some recordings, we computed the oscillation frequency from responses to a full-field step in light intensity, where illumination changed alternatingly from background to white (+100% contrast) or to black (−100% contrast), each for 1 s (Figure S4D). For each trial, we subtracted the average membrane potential measured over the 200 ms prior to the light step and computed the average response trace over trials. We applied an analysis window of 800 ms, ranging from 250 ms after stimulus onset to 50 ms after the end of the contrast step and used the mean-subtracted average response to the preferred contrast. The oscillation frequency was determined as the frequency with maximum power, as assessed by Fourier analysis of the response trace in the analysis window.

### Statistical tests and correlation coefficients

Statistical tests for significance were performed with a Wilcoxon rank sum test (ranksum function in MATLAB) when samples were independent and with a Wilcoxon signed rank test (signrank function in MATLAB) for paired samples. Correlation coefficients between two variables were computed as the Pearson correlation coefficient (corrcoef function in MATLAB).

## Data availability

The data of this study are available at: [*site released upon publication*]

## Author contributions

Conceptualization, H.M.S and T.G.; Methodology, H.M.S and T.G.; Investigation, H.M.S.; Formal Analysis, H.M.S.; Writing, H.M.S. and T.G.; Supervision and Resources, T.G.

## Acknowledgments

The authors would like to thank Michael Weick and Mohammad Khani for their experimental and technical assistance, Mohammad Khani for his help with morphological reconstruction of bipolar cells and fruitful discussions, and Maarten Kamermans and Lauw Klaassen as well as Jochen Staiger and Patricia Sprych for sharing protocols for staining neurons with neurobiotin. This work was supported by the European Research Council (ERC) under the European Union’s Horizon 2020 research and innovation programme (grant agreement number 724822) and by the Deutsche Forschungsgemeinschaft (DFG, German Research Foundation) – Projektnummer 154113120 – SFB 889, project C1.

## Supplemental Information

**Figure S1.**
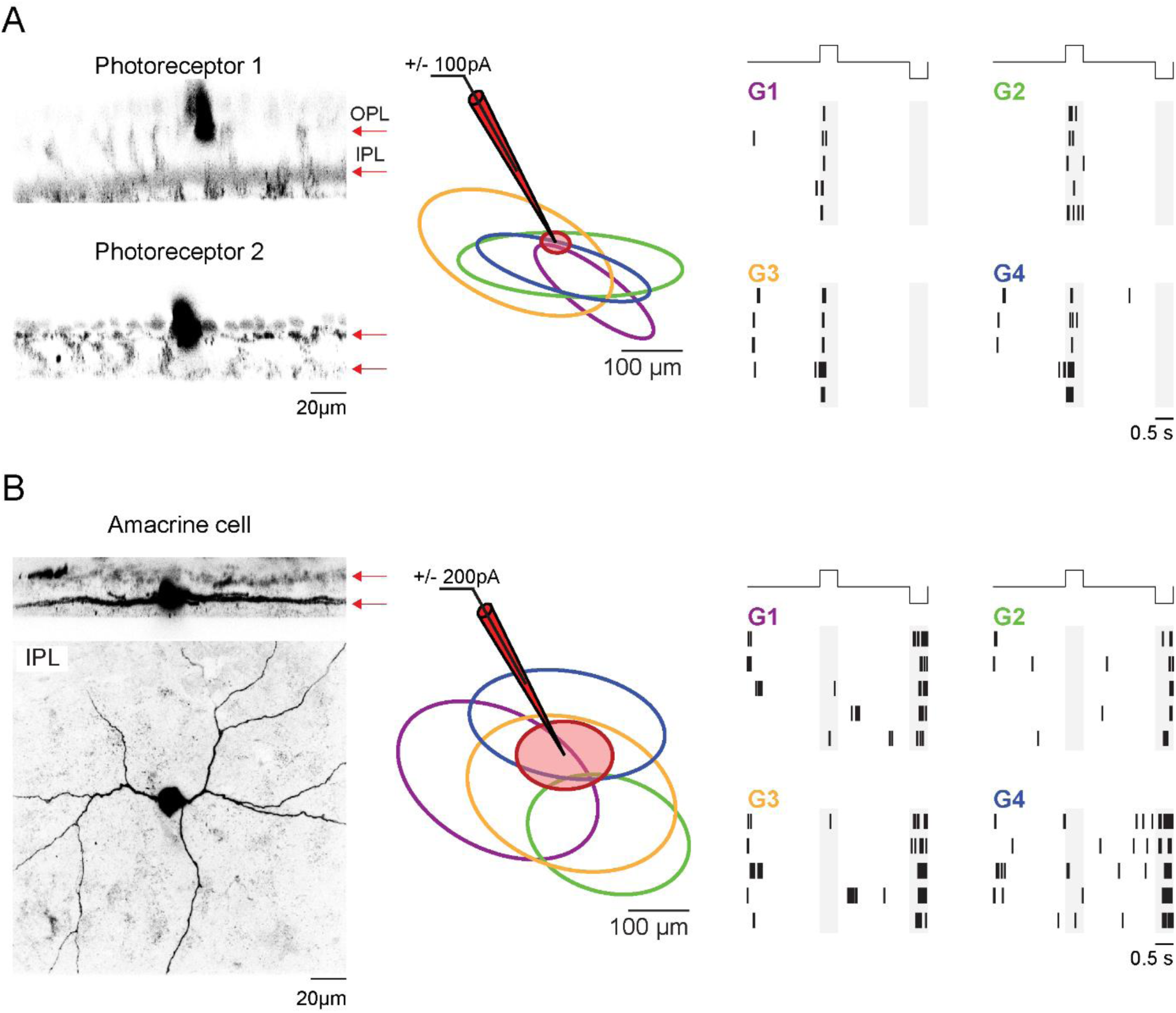
Distinguishing bipolar cells from photoreceptors and amacrine cells. A. Left: Two stained photoreceptors (vertical-section view as in Figure 1B; red arrows mark outer and inner plexiform layers, OPL and IPL). Right: Receptive field outlines and responses of four retinal ganglion cells (color-coded) to current injection into Photoreceptor 1 (receptive field shown in red). All four ganglion cells were OFF cells. B. Left: Stained amacrine cell (vertical- and horizontal-section view in the inner plexiform layer, as in Figure 1B). Right: Receptive field outlines and responses of four ganglion cells to current injection into the amacrine cell. All four cells were OFF cells and responded to negative current injection.

**Figure S2.**
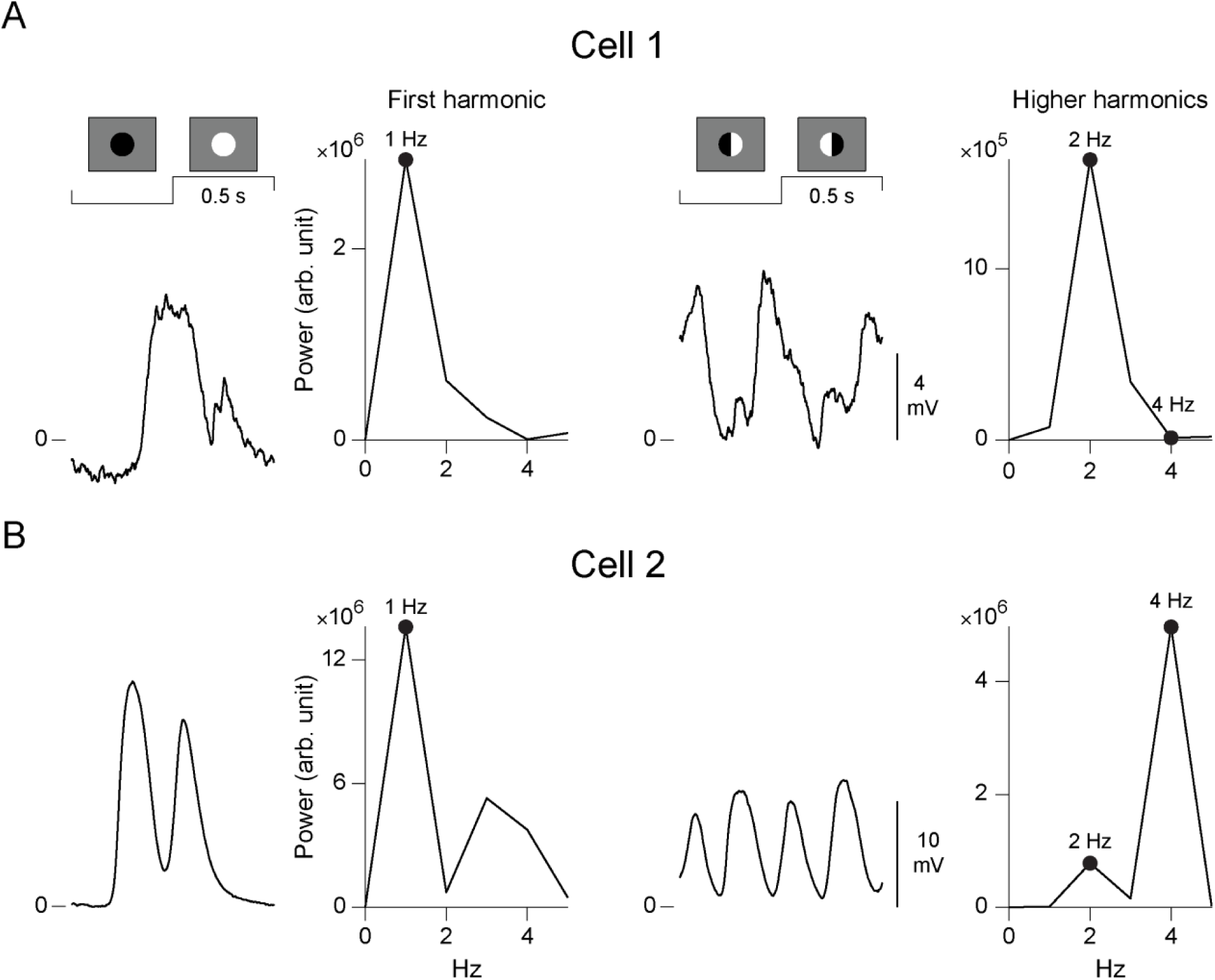
Fourier analysis for computing the spatial nonlinearity index (SNi). A. Left: Trial-averaged response to the uniform contrast-reversing spot and the corresponding power spectrum of the response for one sample cell. The power at the first harmonic, 1 Hz, is marked with a black dot. Right: Trial-averaged response of the same cell to the split spot and the corresponding power spectrum. The black dots mark the power at higher harmonics (2 and 4 Hz). B. Same as A for another sample cell. Note the four peaks in the response trace to the split spot (right side) for Cell 2 and the corresponding peak in the power spectrum at 4 Hz.

**Figure S3.**
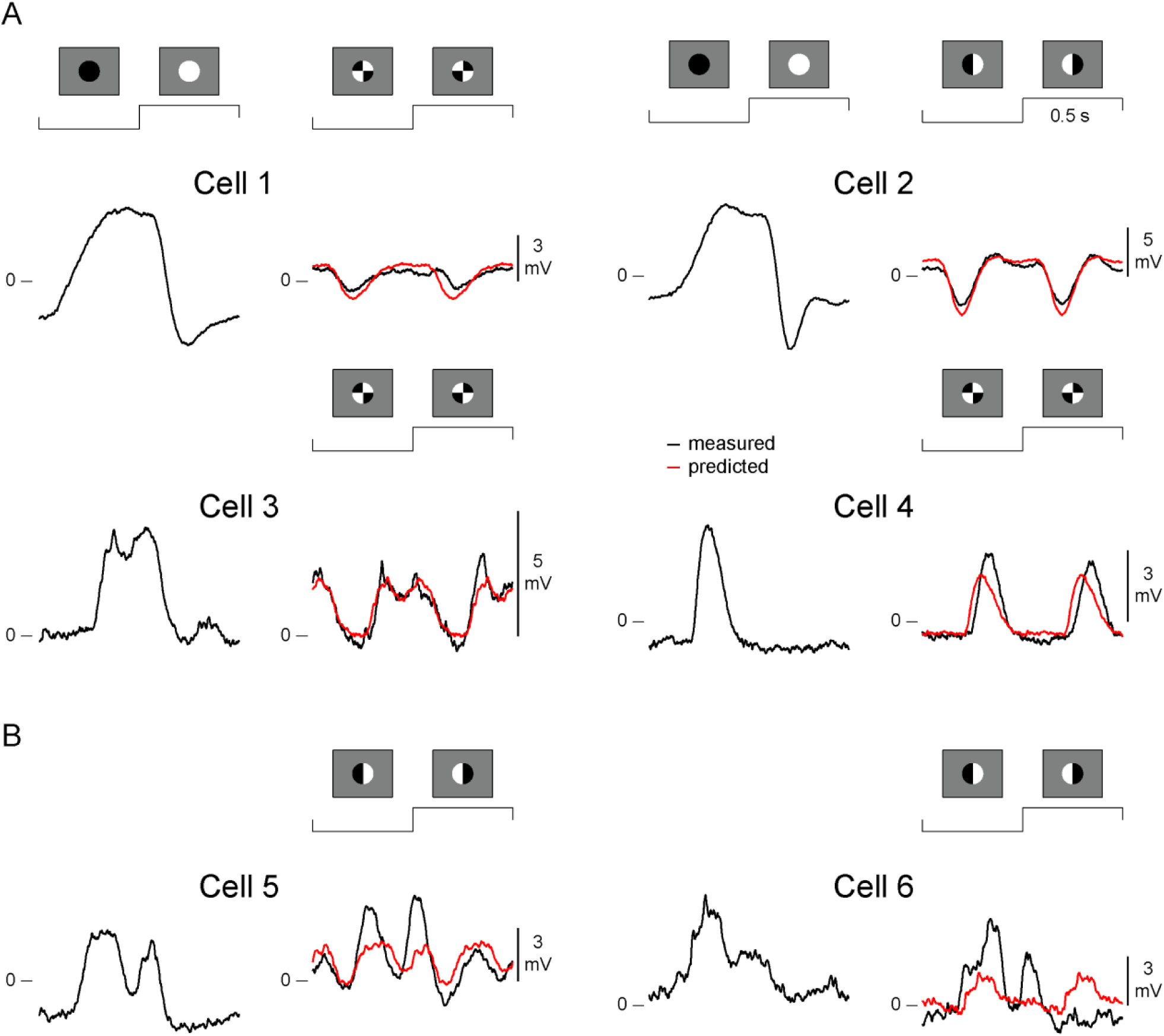
Predicting bipolar cell responses to the patterned-spot presentation. A. Trial-averaged response traces under the uniform contrast-reversing spot and under a patterned spot for 4 sample cells. The prediction to the patterned spot (cf. Figure 6) is shown in red. The prediction accuracy for all four cells is above 0.7. B. Same as A for two sample cells where the prediction accuracy is lower (0.59 for Cell 5 and −0.08 for Cell 6). Note that these two cells responded more strongly to the first reversal of the split spot than to the second, which is an indication that the split spot was not well placed to subdivide the receptive field into equally effective subfields. This may explain the lower prediction accuracy for these cells.

**Figure S4.**
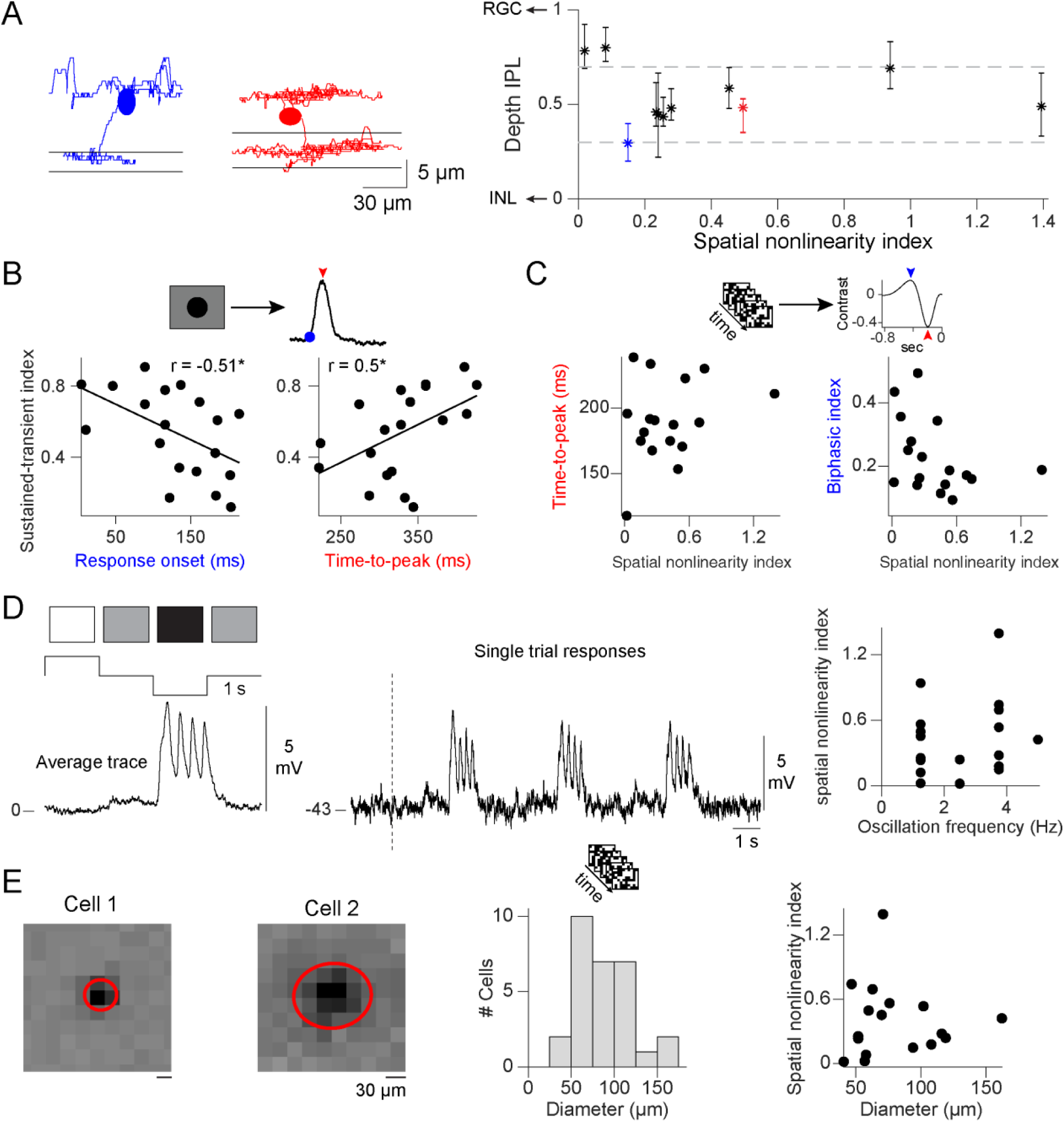
Relation between nonlinear processing and standard response properties in bipolar cells. A. Left: Morphology of two reconstructed bipolar cells. Black lines mark the boundaries of the inner plexiform layer (IPL). Right: Stratification depth within the IPL. Data points mark the mean stratification level of axonal arborization of individual bipolar cells, the error bars the 25th and 75th percentile (n=11). Unity corresponds to the ganglion cell layer and zero to the inner nuclear layer. The data for the two cells depicted on the left are marked by corresponding colors. B. Left: Scatter plots of the sustained-transient index (STi) versus response onset (left; r=-0.51, p=0.026, n=19) and versus time-to-peak (right; r=0.5, p=0.03, n=19), all computed from responses to the local spot of preferred contrast (cf. Figure 2A). Each black dot marks one bipolar cell, and black lines mark the linear regression. C. Same as B for the properties of the temporal filter versus the spatial nonlinearity index. No correlation was observed between the spatial nonlinearity index (SNi) and the time-to-peak of the filter (r=0.26, p=0.3, n=17; left) or between the SNi and the biphasic index (r=-0.39, p=0.13, n=17; right). D. Left: Trial-averaged response trace to full-field contrast light steps that change from gray to white or black in 1-s steps. Note the oscillatory response of the sample cell to the preferred black contrast (∼4 Hz). Middle: Single-trial responses of the same cell. Black dashed line marks the onset of positive contrast (“white”). Right: Scatter plot of oscillation frequency versus the SNi. No correlation was observed (r=0.2, p=0.4, n=19). E. Spatial receptive field components measured with spatiotemporal white noise for two sample bipolar cells and the fitted receptive field outlines (red ellipses). Middle: Distribution of receptive field diameters measured with white noise (range: 41-164 µm, n=29). Right: No correlation was observed between the receptive field diameter and the SNi (r=-0.03, p=0.92, n=17). (*p<0.05)

